# Interleukin-13 and its receptor are synaptic proteins involved in plasticity and neuroprotection

**DOI:** 10.1101/2021.12.01.470746

**Authors:** Shun Li, Florian olde Heuvel, Rida Rehman, Zhenghui Li, Oumayma Aousji, Albrecht Froehlich, Wanhong Zhang, Alison Conquest, Sarah Woelfle, Michael Schoen, Caitlin O’Meara, Richard Lee Reinhardt, David Voehringer, Jan Kassubek, Albert Ludolph, Markus Huber-Lang, Bernd Knöll, Maria Cristina Morganti-Kossmann, Tobias Boeckers, Francesco Roselli

## Abstract

Immune system molecules are expressed by neurons, often for unknown functions. We have identified IL-13 and its receptor IL-13Ra1 as neuronal, synaptic proteins in mouse, rat, and human brains, whose engagement upregulates the phosphorylation of NMDAR and AMPAR subunits and, in turn, increases synaptic activity and CREB-mediated transcription. We demonstrate that increased IL-13 is a hallmark of traumatic brain injury (TBI) in mice as well as in two distinct cohorts of human patients. We also provide evidence that IL-13 upregulation protects neurons from excitotoxic death. We show IL-13 upregulation occurring in several cohorts of human brain samples and in CSF. Thus, IL-13 is a previously unrecognized physiological modulator of synaptic physiology of neuronal origin, with implications for the establishment of synaptic plasticity and the survival of neurons under injury conditions. Furthermore, we suggest that the neuroprotection afforded through the upregulation of IL-13 represents a new entry point for interventions in the pathophysiology of TBI.

## Introduction

Continuous remodelling of synapses at structural and functional level is critical not only to the formation and retention of new memories and to the learning process, but also to the acute and long-term response to pathological conditions. Several components of the innate (Nguyen et al., 2020; Stellwagen and Malenka, 2006; Nelson et al., 2012) and adaptive (Datwani et al., 2009) immune systems have surprisingly been shown to be involved in synaptic plasticity (Boulanger, 2009), either independently of their primary immune function (Bialas and Stevens, 2013; Datwani et al., 2009; Stellwagen and Malenka, 2006; Wang et al., 2021) or as a consequence of the modulation of local microglia (Nguyen et al., 2020). The source of many of these immune-related mediators appears not to be infiltrating immune cells but, rather, glial cells such as astrocytes (Stellwagen and Malenka, 2006; Wang et al., 2021) and even the neurons themselves (Gruol, 2016; Olde Heuvel et al., 2019). Nevertheless, synapses may be sensitive not only to cytokines produced by CNS cells but also to the large amounts secreted by infiltrating inflammatory cells in autoimmune diseases (Centonze et al., 2009) or in TBI (Morganti-Kossmann et al., 2019).

Recent evidence (Olde Heuvel et al., 2019; Mori et al., 2017) suggests that IL-13, besides its immune functions, may also have a neuronal origin as well as neuromodulatory properties, which may affect spatial learning (Brombacher et al., 2017). IL-13 shares approximately 30% of homology with IL-4 (Zurawski and de Vries, 1994), but is strongly related to the latter in their ability to enhance immune responses driven by subsets of Th2 lymphocytes and in the IgE switch in B lymphocytes (McKenzie, 2000). The immune sources of IL-13 include mast cells, basophils and eosinophils, as well as CD4 T cells, ILC2 and NK T cells (Juntilla, 2018). IL-13 signals through a type-I receptor, composed by the specific IL13-Ra1 subunit that, upon ligand binding, forms a heterodimer with IL-4 receptor alpha chain and drives a JAK2/Tyk2-dependent phosphorylation of STAT6, activating transcriptional responses (Goenka and Kaplan, 2011). The type-II receptor, the high-affinity IL-13Ra2, does not have a cytoplasmic chain and, although originally considered a decoy, appears to activate a number of signaling cascades (Fichter-Feigel et al., 2006).

IL-13 is an effector cytokine, acting upon epithelial cells and smooth muscle cells in the gut and airways and triggering itch in the skin (Roesner et al., 2019); as such, IL-13 has been extensively studied in the context of the pathogenesis of allergic diseases such as asthma (Wills-Karp et al., 1998) and atopic dermatitis (Furue et al., 2019). In fact, blockade of peripheral IL-13 by monoclonal antibodies has emerged as a promising therapeutic option in these conditions (Corren et al., 2011; Rabe et al., 2018).

In this study, we have investigated the biology of neuronal IL-13. We demonstrate that IL-13 and IL-13Ra1 are synaptic components expressed in rat, mouse and human neurons in an activity-dependent manner in normal conditions and, most notably, that IL-13 is upregulated upon traumatic injury. Furthermore, we show that IL-13 triggers the phosphorylation of glutamate receptors and several presynaptic proteins; it increases synaptic activity and neuronal firing ultimately driving the phosphorylation of several transcription factors, including CREB. Finally, we reveal that IL-13 protects neurons against excitotoxic insults, implying a direct neuroprotective role in TBI. We use human samples of brain and CSF from three distinct normal and TBI cohorts to demonstrate the relevance of IL-13 in the physiopathology of TBI in patients.

## Results

### 1. IL13 and its receptor IL-13Ra1 are neuronal synaptic proteins

We set out to establish the source(s) and the localization of IL-13 and its receptor in the cerebral cortex of the mouse. First, we explored the expression of IL-13 using *in situ* hybridization. Surprisingly, we found that a substantial fraction of cells expressed abundant levels of IL-13 mRNA; double *in situ* hybridization using markers of glutamatergic neurons (VGLUT1 for layer II/III, VGLUT2 for layer IV; Fremeau et al., 2001, Figure 1A and Supplementary Figure 1A-B) and GABAergic neurons (VGAT, Figure 1B) revealed that IL-13 is expressed in both cell subpopulations. When the overall cortical population was used as reference, we found that VGLUT1+ glutamatergic neurons expressed higher levels of IL-13 than the global cell population, whereas GABAergic neurons expressed lower levels of IL-13 than the global cell population, implying that IL-13 was more strongly expressed in excitatory neurons (Figure 1C). Interestingly, IL-13 was also expressed in VGLUT2+ Layer IV neurons, which displayed average expression levels comparable to VGLUT1+ neurons. Notably, a substantial degree of variability in expression levels was observed across glutamatergic neurons (Supplementary Figure A-B) pointing toward a regulated, rather than constitutive, expression.

**Figure 1:**
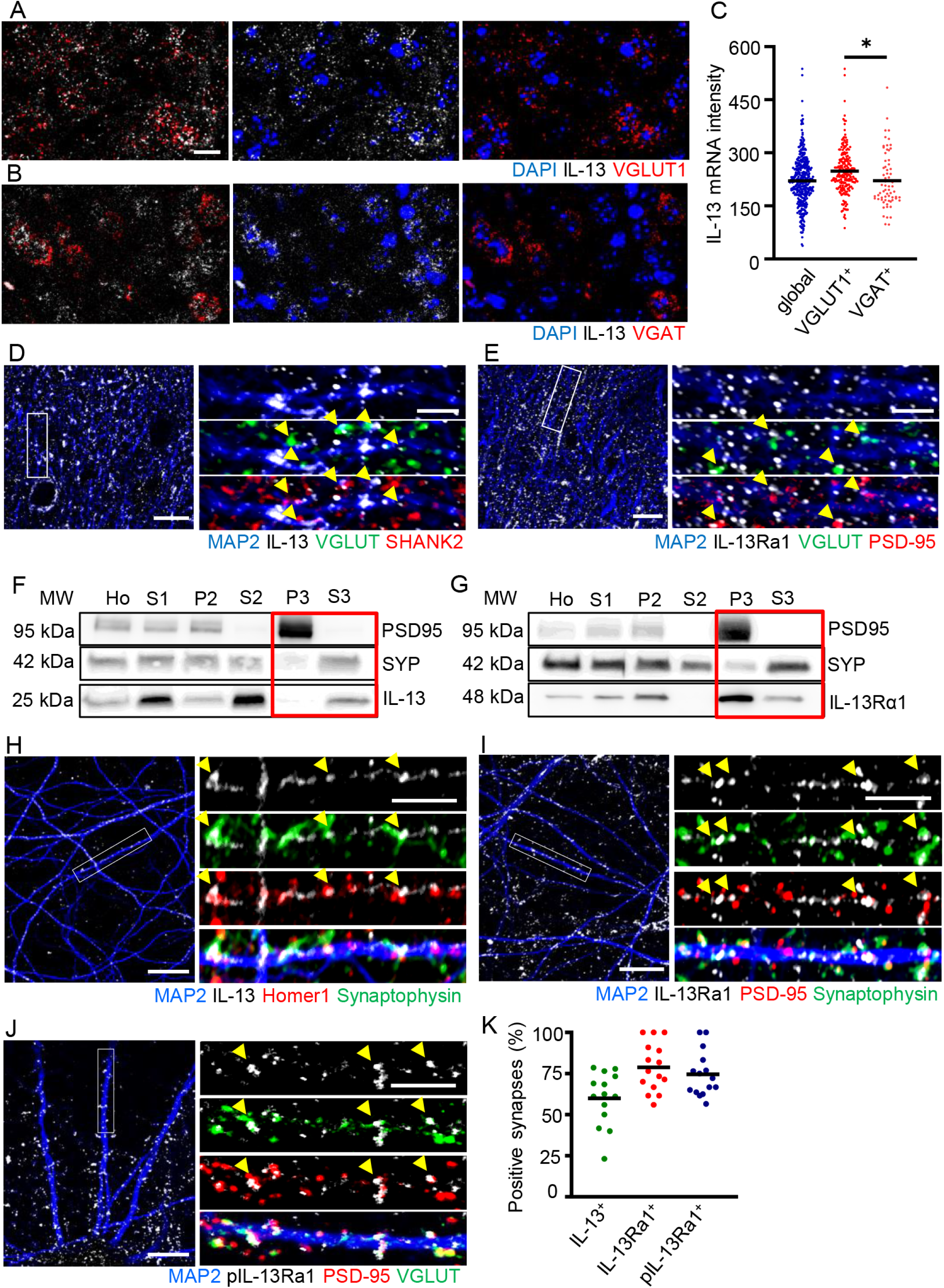
Neuronal IL-13 and its receptor IL-13Ra1 are synaptic proteins. A. IL-13 mRNA expression in VGLUT1 positive glutamatergic neurons in layer II/III of mouse cortical sections (single molecule in situ hybridisation). N = 3. Scale bar: 20μm. B. IL-13 mRNA expression in VGAT positive GABAergic neurons in layer II/III of mouse cortical sections (single molecule in situ hybridisation). N = 3. Scale bar: 20μm. C. Increased IL-13 mRNA intensity in VGLUT1 positive population compared to VGAT positive and global cell populations in layer II/III of mouse cortical sections. N = 3. * : p<0.05. D. Synaptic localisation of IL-13 in mouse cortical sections (Immunostaining with MAP2, pan-VGLUT and SHANK2). N = 3. Scale bar overview 10μm and insert 5μm. E. Synaptic localisation of IL-13Ra1 in mouse cortical sections (Immunostaining with MAP2, pan-VGLUT and PSD-95). N = 3. Scale bar overview 10μm and insert 5μm. F. Fractionation experiment in mouse cortical tissue show IL-13 located in homogenates (Ho), homogenates without nuclei, cell debris and extracellular matrix (S1), crude membrane fraction (P2), cytosolic compartment (S2) and mainly in the presynaptic cytosol (S3), but almost none in the post-synaptic density (P3). verified by blotting with PSD-95 and synaptophysin. N = 3. G. Fractionation experiment in mouse cortical tissue show IL-13Ra1 located in homogenates (Ho), homogenates without nuclei, cell debris and extracellular matrix (S1), crude membrane fraction (P2), cytosolic compartment (S2) and mainly in the post-synaptic density (P3), with a small fraction in the presynaptic cytosol fraction (S3). verified by blotting with PSD-95 and synaptophysin. N = 3. H. Synaptic localisation of IL-13 in rat cortical neurons (Immunostaining with MAP2, synaptophysin and Homer1). N = 3. Scale bar overview 10μm and insert 5μm. I. Synaptic localisation of IL-13Ra1 in rat cortical neurons (Immunostaining with MAP2, synaptophysin and PSD-95). N = 3. Scale bar overview 10μm and insert 5μm. J. Synaptic localisation of pIL-13Ra1 in rat cortical neurons (Immunostaining with MAP2, pan-VGLUT and PSD-95). N = 3 Scale bar overview 10μm and insert 5μm. K. IL-13, IL-13Ra1 and pIL-13Ra1 show 60%, 79% and 75%, respectively, of colocalization with mature synapses. N = 3.

Secondly, we further investigated the localization of IL-13 protein and its cognate receptor IL-13Ra1 in the cortex by immunostaining. In murine cortex, the synapses were identified as areas showing colocalization of the pre- and post-synaptic markers such as VGLUT+/SHANK+ or VGLUT/PSD-95+, along MAP2+ dendrites. Approximately 50% of bona fide synapses showed immunoreactivity for IL-13 (43±9%, Figure 1D) or, in independent experiments, for IL-13Ra1 (47±9%, Figure 1E). These findings raised the possibility that IL-13 and IL-13Ra1 are not only neuronal proteins but are actual synaptic proteins. We then sought an independent confirmation of the synaptic nature of IL-13 and IL-13Ra1 using a brain fractionation protocol to isolate distinct cellular subcompartments (Reim D et al., 2017), namely whole cortex extracts, neuronal membranes, synaptosomes, post-synaptic membranes and presynaptic vesicles. The enrichment of the individual fractions was monitored by Western Blot using antibodies against PSD-95 and Synaptophysin as markers of the post-synaptic and the vesicle fractions, respectively. When the protein extract from each fraction was probed with an antibody against IL-13, the immunoreactivity was strongly enriched in the synaptic vesicle fraction (S3) and mostly excluded from the postsynaptic fraction (P3) in a pattern closely matching that of synaptophysin (Figure 1F). Interestingly, when the same fractions were probed with an antibody directed against IL-13Ra1, we identified a very strong enrichment in the post-synaptic fraction, compared with a low representation in the whole-cortex homogenate and the absence in the presynaptic vesicle fraction (thus resembling the pattern of PSD-95, Figure 1G). Thus, these findings not only confirm the synaptic localization of IL-13 and IL-13Ra1 but suggest a pre- and post-synaptic (respectively) enrichment.

Thirdly, since the bulk cortical tissue has a very complex architecture with high density of intertwined astrocytes and neuronal processes, we set out to elucidate the synaptic localization of both IL-13 and IL-13Ra1 in cultured rat cortical neurons. In E18-DIV21 cultures (which contain approx. 6% astrocytes and no microglia) we detected a robust expression of mRNA for IL-13, IL-13Ra1 and IL-13Ra2 (Supplementary Figure 1F). Cultured neurons displayed a large number of synaptic contacts (PSD-95+ and pan-VGLUT+) along with the dendrites (identified by MAP2 immunostaining) and approximately 50% of established synapses (i.e., characterized by the colocalization of pre- and post-synaptic markers) overlapping with IL-13, IL-13Ra1 or pIL-13Ra1 (Figure 1H-K). Colocalization was higher in those synapses that were positive for both markers rather than for one or the other, indicating that mature synapses are more prominently enriched in IL-13/IL-13Ra1. To minimize the risk of any unpredicted and unspecific staining generated by anti-IL-13 mouse monoclonal antibody, we repeated this set of experiments with an unrelated rabbit polyclonal antibody against IL-13. The findings obtained with the second antibody closely overlapped with the patterns detected with the mouse monoclonal (Supplementary Figure 1E). We further validated the reagents used for the immunolocalization of IL-13 using brain tissue from IL-13^-/-^ mice (McKenzie et al., 1998): both the mouse and the rabbit antibodies generated negligible staining in the IL-13^-/-^ samples (whereas immunoreactivity was identified in their WT littermates; Supplementary Figure 2A-B).

### 2. Super-resolution microscopy reveals pre- and post-synaptic localization of IL-13 and IL-13Ra1

Since the resolution of the diffraction-limited confocal microscopy prevented the establishment of pre- or post-synaptic nature for IL-13 and its receptor, we opted to use a STED nanoscopic imaging paradigm to discern their localization. Cultured cortical neurons were immunostained for PSD-95 (post-synaptic marker) or Bassoon (pre-synaptic marker) together with IL-13 or IL-13Ra1. We plotted the relative distribution of the peaks of IL-13 relative to either PSD-95 or Bassoon (using the MAP2 profiles to establish the synaptic polarity). Most notably, the peak of IL-13 distribution was located approximately 150nm away from the PSD-95 peak (n=200 synapses), with a tail extending up to 300 nm away from the dendritic side (Figure 2A and E). On the contrary, IL-13 peak largely overlapped with the Bassoon peak (n=200 synapses), with some of the IL-13 immunoreactivity being localized further away in the presynaptic terminal (Figure 2C and F). These findings, in remarkable agreement, corroborate the presynaptic nature of IL-13. In neurons immunostained for IL-13Ra1, on the other hand, the distribution appeared broader but the peak of IL-13Ra1 was localized within 50nm of the PSD-95 peak on the post-synaptic side (as determined using the MAP2+ profile of the dendrite; n=200 synapses; Figure 2B and E); a close inspection of the STED images revealed that often multiple IL-13Ra1 clusters were localized within the PSD-95+ cluster (Supplementary Figure 1G). In contrast, within samples immunostained for IL-13Ra1 and Bassoon, the IL-13Ra1 peak was localized approximately 200nm away (toward the dendrite) from the Bassoon peak (Figure 2D and F); occasionally we detected smaller IL-13Ra1 located around the Basson spots. Thus, the STED imaging data provide strong evidence for a presynaptic localization of IL-13 and a (mainly) postsynaptic localization of IL-13Ra1 in cultured neurons, in remarkable agreement with the fractionation data from the bulk brain. However, the existence of a smaller, presynaptic pool of IL-13Ra1 cannot be currently discounted.

**Figure 2:**
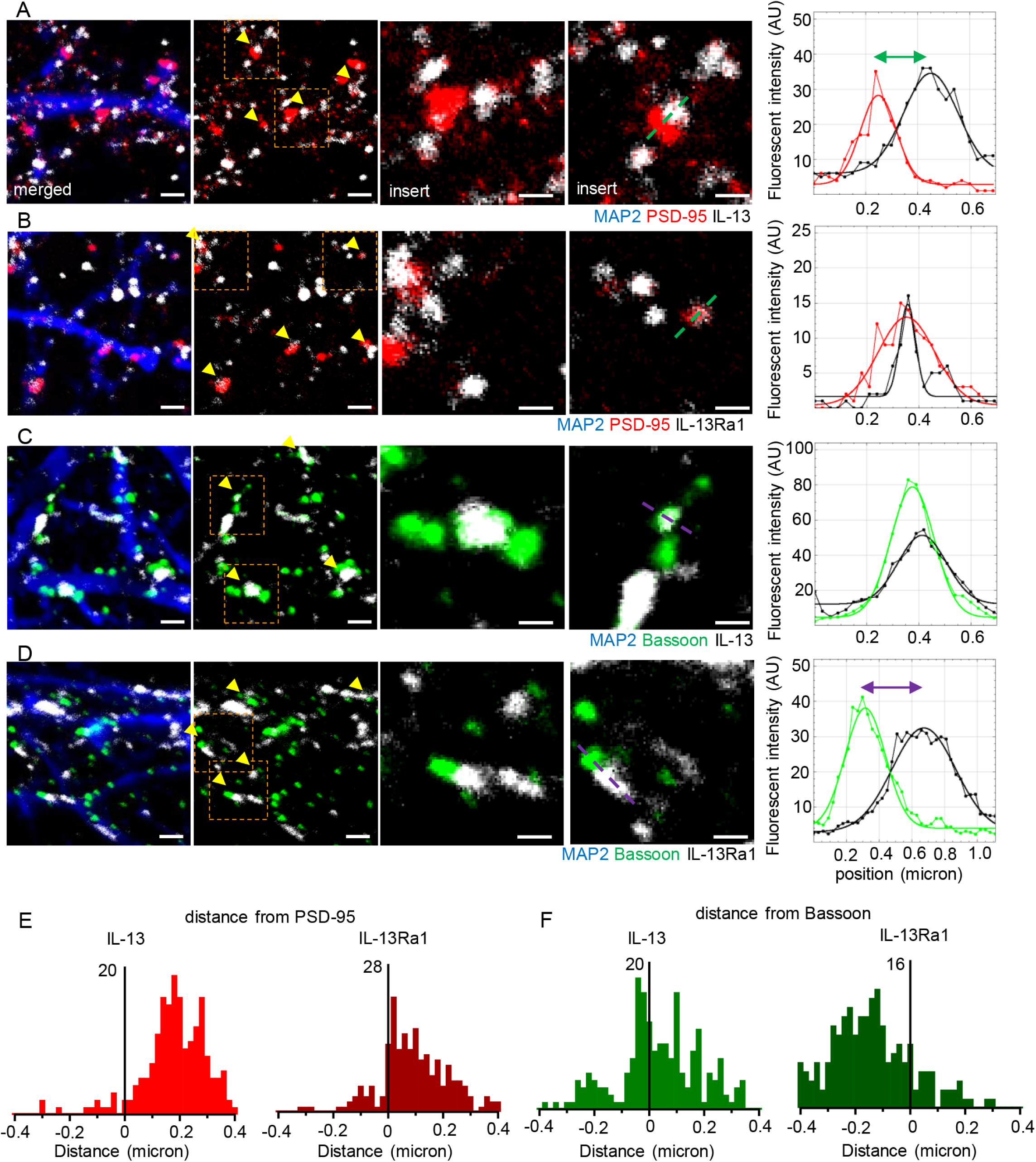
Presynaptic IL-13 and a postsynaptic lL-13Ra1 demonstrated by super-resolution microscopy. A. STED images and intensity profiles plots of single synapses show a distinct separation between PSD-95 and IL-13. N = 200 synapses. Scale bar overview: 1μm, scale bar insert: 500nm. B. STED imaging and intensity profile plots of single synapses show an overlap between PSD-95 and IL-13Ra1. N = 200 synapses. Scale bar overview: 1μm, scale bar insert: 500nm. C. STED imaging and intensity profile plots of single synapses show an overlap between Bassoon and IL-13. N = 200 synapses. Scale bar overview: 1μm, scale bar insert: 500nm. D. STED imaging and intensity profile plots of single synapses show a distinct separation between Bassoon and IL-13Ra1. N = 200 synapses. Scale bar overview: 1μm, scale bar insert: 500nm. E. Analysis of distance between peaks reveal a distance between IL-13 and PSD-95 with two distinct peaks, one at 150 nm and the second at 300 nm. Analysis of distance between peaks reveal a distance between IL-13Ra1 and PSD-95 with a broader distribution of IL-13Ra1, but a main peak close to PSD95. N = 200 synapses. F. Analysis of distance between peaks reveal a distance between IL-13 and Bassoon with multiple distinct peaks, one at overlapping Bassoon and some further away (100 and 200 nm). Analysis of distance between peaks reveal a distance between IL-13Ra1 and Bassoon with a broader distribution of IL-13Ra1, but a main peak at -150 nm from Bassoon. N = 200 synapses.

Taken together (Figure 1 and Figure 2), this data identifies IL-13 and IL-13Ra1 as neuronal, synaptic proteins in rat and mouse brain and indicate a polarized biology with IL-13 released by presynaptic terminals acting on post-synaptic IL-13Ra1 receptors.

### 3. IL-13 triggers a large-scale phosphorylation of glutamate receptors and presynaptic proteins

If IL-13 and its receptor were previously not recognized as synaptic proteins, what is their fucntion in synaptic biology? To gain insights into the roles of synaptic IL-13, we used phospho-antibody arrays to characterize the large-scale architecture of phosphorylation events in neurons set in motion by the cytokine. We treated cultured cortical neurons with IL-13 (or vehicle) for 1h or 3h; whole-cells protein extract was then processed using a glass-based phospho-antibody array assay involving multiple phosphorylation sites on 167 distinct targets (including ion channels, neurotransmitter receptors, vesicle proteins and cytoskeletal elements, among the others).

At 1h post stimulation, 24/167 proteins displayed an increase in phosphorylation, whereas 15/167 proteins were down-phosphorylated on several serine, threonine or tyrosine phospho epitopes. Intriguingly, both pre- and post-synaptic proteins were significantly represented in the hit list. Among the post-synaptic proteins, IL-13 increased the phosphorylation of NMDAR1 (S897), of AMPAR1 (GluR1, both S849 and S863), of CaMKII (T286 and T305) and, interestingly, of the BDNF receptor TrKB (Y515 and Y705). Among the presynaptic proteins, IL-13 induced the phosphorylation of α-synuclein (Y133 and Y136) and synapsin I (S62). A substantial number of signaling cascades was also activated, including CaMKI and CaMKIV (S177 and S196/200, respectively), GSK3β (Y216/279), PP2A (Y307), PKA (T197) and GRK2 (S685). Other notable proteins up-phosphorylated by IL-13 included cytoskeletal proteins Tau (S231) and Merlin (S518) (Figure 3A-B). Among the down-regulated phospho-epitopes, were TrkA (Y680, Y701 and Y791) and TrkC (Y516) and the GAB family members GAB1 and GAB2 (Y627 and Y643, respectively); also the NMDAR1 epitope S896 showed a degree of downregulation (Figure 3A-B).

**Figure 3:**
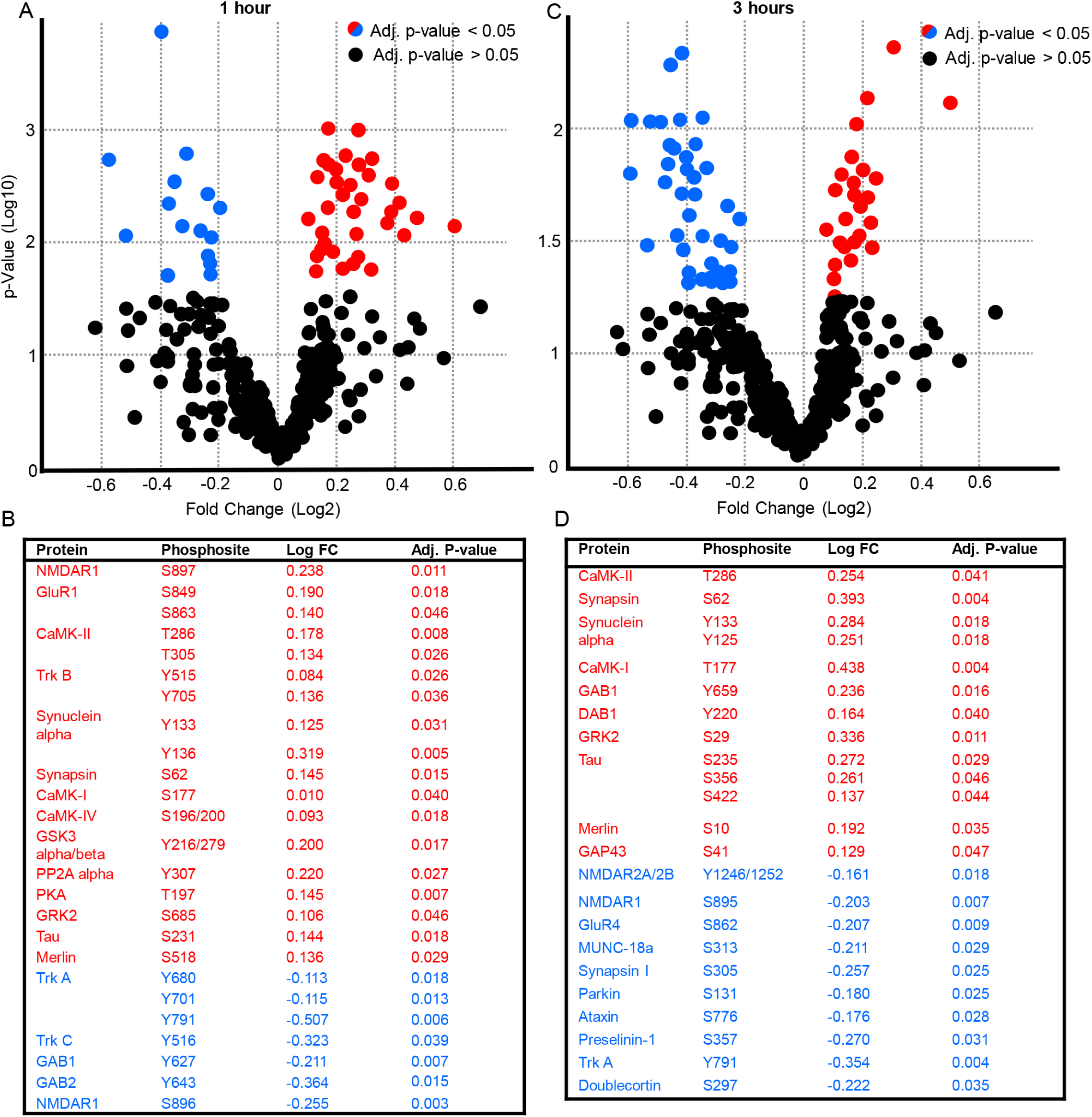
IL-13 causes the large-scale phosphorylation of glutamate receptors and presynaptic proteins. A-B. Volcanoplot and list of proteins showing significant change in their phosphorylation (up or down) 1 hour after neurons were exposed to IL-13 (50 ng/ml). red = up-phosphorylated, blue = down-phosphorylated. N = 4. C-D. Volcanoplot and list of proteins showing significant change in their phosphorylation (up or down) 3 hour after neurons were exposed to IL-13 (50 ng/ml). red = up-phosphorylated, blue = down-phosphorylated. N = 4. Complete list of significantly phosphorylated proteins can be found in supplementary information.

The phosphorylation landscape at 3h post stimulation displayed some degree of similarity with the 1h, but also substantial differences. Overall, 19 proteins (out of 167 included in the array) were up-phosphorylated and 21 were down-phosphorylated. Again, both pre- and post-synaptic proteins were represented among the up-phosphorylated. Among the post-synaptic proteins, only CaMKII (T286) was still significantly up-phosphorylated. Among the pre-synaptic proteins, Synapsin I (S62) and α-synuclein (Y125 and Y133) were up-phosphorylated. A number of additional signaling proteins were still substantially up-phosphorylated, such as CaMK1 (T177), GAB1 (Y659), DAB1 (Y220) and GRK2 (S29). Moreover, the cytoskeletal protein Tau (S235, S356, S422), Merlin (S10) and GAP43 (S41) were still strongly phosphorylated (Figure 3C-D). However, glutamate receptors were prominently represented among the proteins with decreased phosphorylation: NMDAR2A/2B (Y1246/1252), NMDAR1 (S895), AMPAR GluR4 (S862). Two pre-synaptic proteins appeared among the down-phosphorylated proteins, namely MUNC-18a (S313) and Synapsin I (S605). Several down-phosphorylated proteins were involved in protein degradation or post-translational processing, such as Parkin 1 (S131), Ataxin 1 (S776) and Preselinin-1 (S357). Other notable proteins displaying a reduced phosphorylation included TrkA (Y791) and Doublecortin (S297) (Figure 3C-D).

Thus, IL-13 induced the rapid (1h) phosphorylation of NMDA and AMPA-type glutamate receptors (mainly at sites associated with increased trafficking to synapse and membrane insertion) and signaling proteins associated with synaptic plasticity (such as CaMKII and PKA), together with several presynaptic and cytoskeletal proteins. However, the phosphorylation of glutamate receptors was short-lived and at 3h most glutamate receptors were down-phosphorylated (indicating removal from synaptic sites), although signaling associated with CaMKII and presynaptic modifications were still persisting.

### 4. IL-13 drives the phosphorylation of CREB and the induction of activity-dependent immediate-early genes

The pattern of increased phosphorylation of glutamate receptor phosphorylation and CamKII suggests that IL-13 may activate transcriptional programs associated with neuronal activity.

As a preliminary step, we verified the activation of IL-13-triggered signaling cascades in neurons using experiments independent of antibody arrays. We immunostained cultured cortical neurons treated with IL-13 for 1h and 3h for phosphorylated IL-13Ra1 (Tyr405) and for phospho-ERK1/2 (Thr202/Tyr204). Compared to baseline, IL-13 triggered a significant elevation in the phosphorylation of IL-13Ra1 at 1h and, albeit reduced in magnitude at 3h as well (Figure 4A-B and Supplementary Figure 4A-D). Likewise, IL-13 triggered a strong elevation in ERK phosphorylation at 1h, which subsumed by 3h (Figure 4C-D).

**Figure 4:**
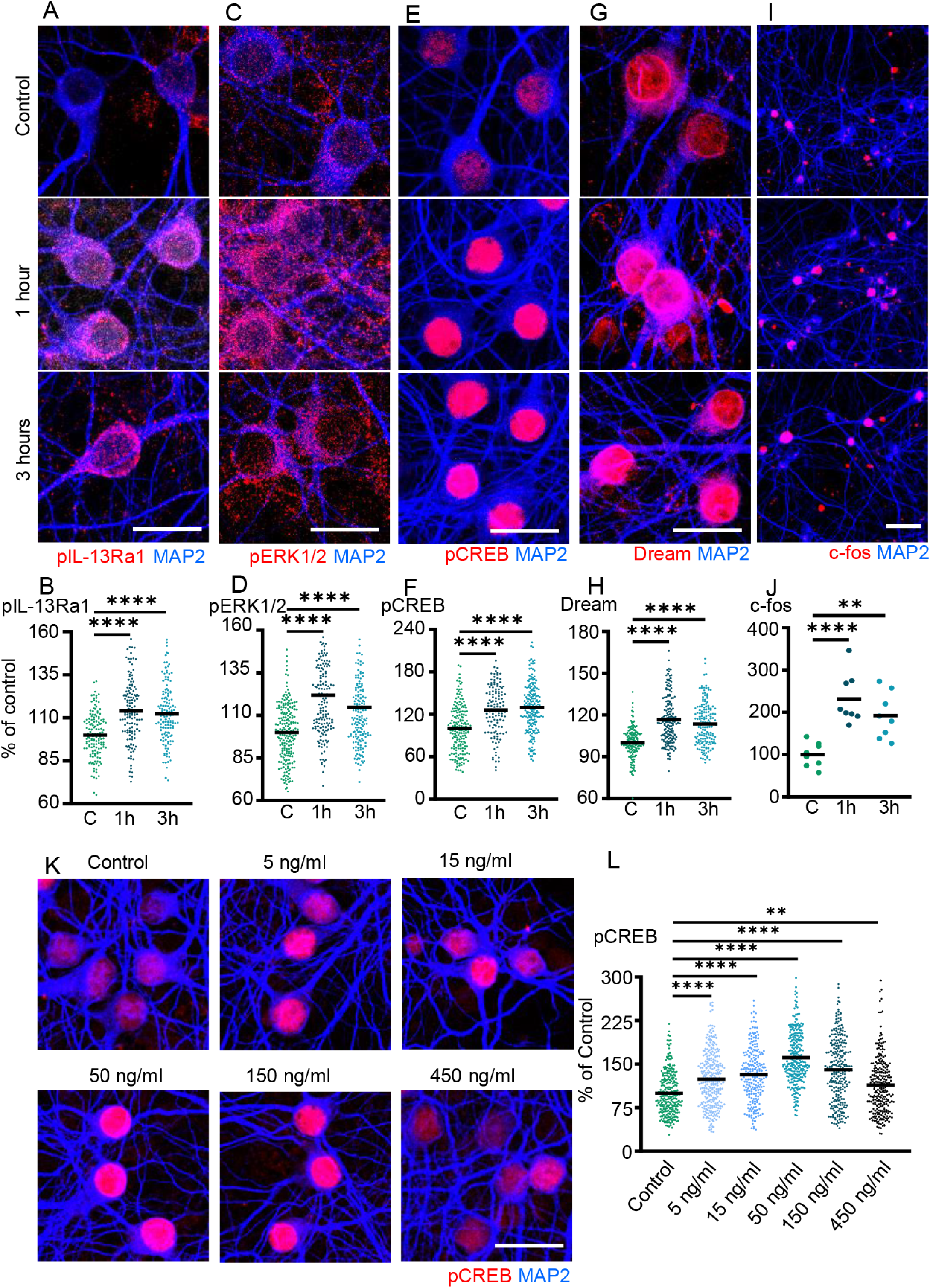
IL-13 induces CREB phosphorylation and immediate-early genes transcription. A-B. Significant up-phosphorylation of IL-13Ra1 in rat cortical neurons 1h and 3h after IL-13 treatment (50 ng/ml) (immunostaining). N = 3. Scale bar: 20μm. ****: p<0.0001 C-D. Significant up-phosphorylation of ERK1/2 in rat cortical neurons 1h and 3h after IL-13 treatment (50 ng/ml) (immunostaining). N = 3. Scale bar: 20μm. ****: p<0.0001. E-F. Significant up-phosphorylation of CREB in rat cortical neurons 1h and 3h after IL-13 treatment (50 ng/ml) (immunostaining). N = 3. Scale bar: 20μm. ****: p<0.0001. G-H.Significant up-phosphorylation of DREAM in rat cortical neurons 1h and 3h after IL-13 treatment (50 ng/ml) (immunostaining). N = 3. Scale bar: 20μm. ****: p<0.0001. I-J. Significant increase of c-fos positive cells in rat cortical neurons 1h and 3h after IL-13 treatment (50 ng/ml) (immunostaining). N = 3. Scale bar: 50μm. **: p<0.01; ****: p<0.0001. K-L. Dose-dependent effect of IL-13 treatment on CREB phosphorylation until 50 ng/ml, doses exceeding this limit reduces phosphorylation of CREB. N = 3. Scale bar: 20μm. **: p<0.01; ****: p<0.0001.

Next, we focused on the phosphorylation of CREB, under the hypothesis that the increased phosphorylation of glutamate receptors upon IL-13 treatment may lead to increased neuronal firing. Indeed, IL-13 treatment caused the massive rise in the levels of phosphorylated CREB (S133) after 1h and 3h (compared to baseline; Figure 4E-F). Most notably, when we treated neurons with ascending concentrations of IL-13 we did not obtain a linear increase in pCREB levels, but rather an inverse-U distribution: effect on pCREB levels increased for IL-13 doses of 5 to 50 ng/ml ( 24%, 32% to 61% of vehicle, respectively) but then declined with doses up to 450 ng/ml (40% of baseline for 150 ng/ml and 14% for 450 ng/ml; Figure 4K-L).

Finally, the elevation in pCREB levels suggest that IL-13 may induce a large-scale transcription of immediate-early genes and other activity-dependent transcription factors. Indeed, we found that IL-13 induced the upregulation of nuclear levels of ATF-3 (Supplementary Figure 3E-F), and DREAM after 1h and 3h treatments (Figure 4G-H), and increased the number of c-fos-positive neurons in the culture (Figure 4I-J). We confirmed this effect by verifying mRNA upregulation of a number of immediate-early genes *(c-fos, fos-B, egr-1, egr-2, gadd45a and gadd45b*) by qPCR (Supplementary figure 3G). In addition, following IL-13 treatment at 1h and 3h we observed the elevation of the phosphorylation of both STAT6 and STAT3 (Supplementary Figure 3C-F), which are well known to be IL-13/IL-13Ra targets (Hershey, 2003).

### 5. IL-13 activates synaptic signaling cascades converging on CREB

Our array screening identified a substantial number of signaling cascades set in motion by IL-13 and several substrates (in particular, several NMDA and AMPA glutamate receptors). In order to i) validate the functional engagement of signaling kinases and ii) determine their involvement in transcriptional responses evoked by IL-13, we performed a small-molecule inhibitor screening, using CREB phosphorylation as readout (being the most robust and well-understood of the TF involved).

IL-13-induced phosphorylation of CREB was completely prevented by the JAK inhibitor Ruxolitinib and by the ERK inhibitor PD98059, in agreement with their role in proximal signaling events upon IL-13Ra1 engagement (Figure 5A-B). Among the downstream signaling molecules identified by the array, we then verified the role of Cdk-5, GSK-3β CaMKII and PKA (using Roscovitine, CHIR, KN-93 and H89); we also studied the effects of a TrkB inhibitor, ANA-12 (since TrkB appears among the up-phosphorylated proteins, Figure 3A-B)). IL-13-induced phosphorylation of CREB was prevented by PKA and CaMKII inhibitors, but not by Cdk5 or GSK-3β inhibitors (Figure 5E-F), underscoring the prominent role of synapse-signaling cascades in driving CREB phosphorylation. Interestingly, ANA-12 partially decreased IL-13 effect, indicating some degree of involvement of BDNF signaling in IL-13 effects.

**Figure 5:**
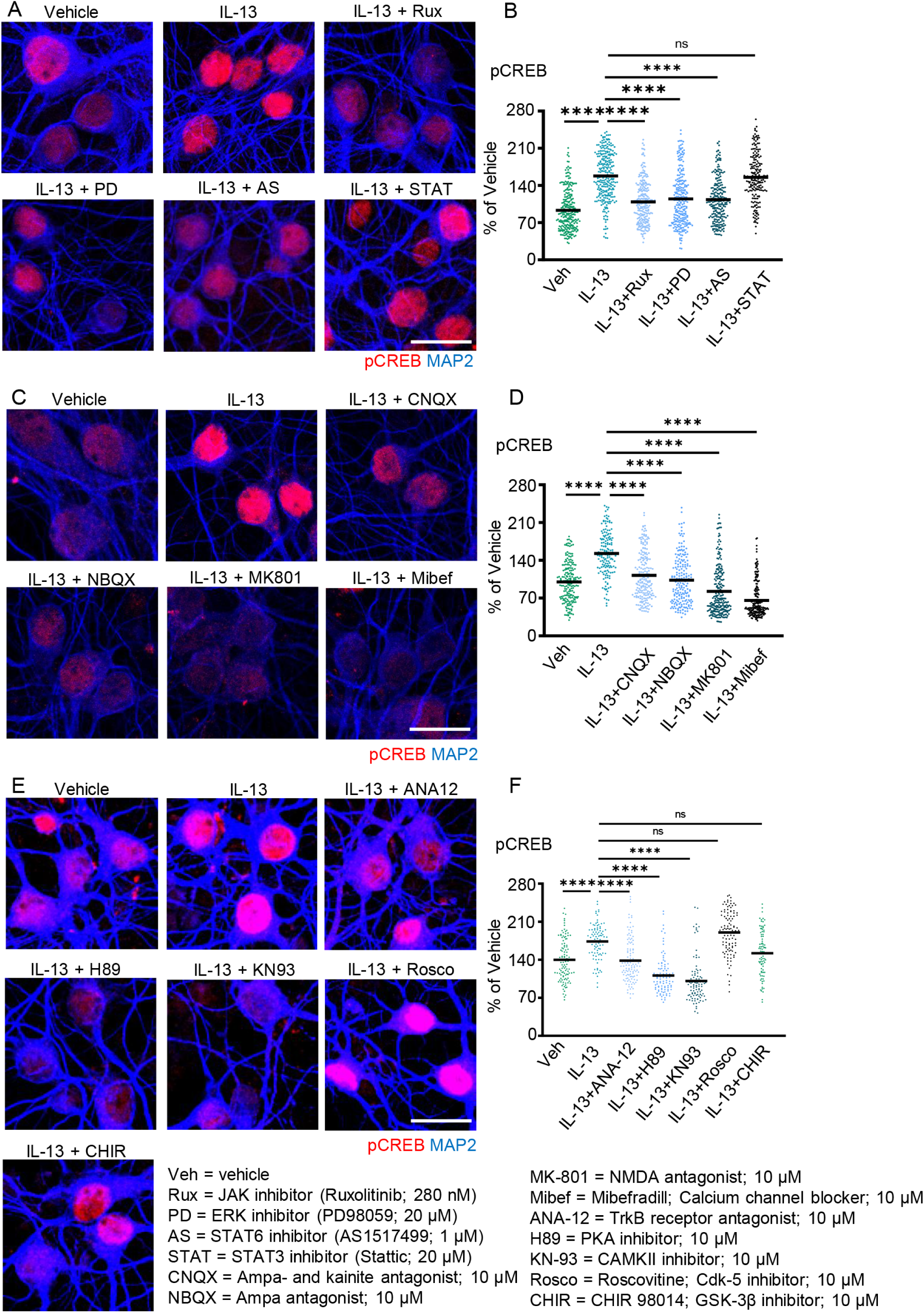
IL-13 activates CREB phosphorylation through JAK/ERK1/2 and NMDAR/AMPAR signaling. A-B. JAK (Ruxolitinib; 280nM), ERK1/2 (PD98059; 20 μM) and STAT6 (AS1517499; 1 μM) inhibitors significantly abolish the IL-13 dependent CREB phosphorylation in rat cortical neurons. STAT3 inhibitor (Stattic; 20 μM) did not alter the IL-13 induced phosphorylation of CREB. N = 3. Scale bar: 20μm. ****: p<0.0001. C-D. AMPA receptor antagonists (CNQX, NBQX; both 10 μM), NMDA receptor antagonist (MK-801; 10 μM) and calcium channel blocker (Mibefradil; 10 μM) significantly reduce the IL-13 induced CREB phosphorylation. N = 3. Scale bar: 20μm. ****: p<0.0001. E-F. TrkB receptor antagonist (ANA12; 10 μM), PKA inhibitor (H89; 10 μM) and CAMK-II inhibitor (KN93; 10 μM) significantly reduce the IL-13 induced CREB phosphorylation. CDK5 inhibitor (Roscovitine; 10 μM) and GSK-3 inhibitor (CHIR 98014; 10 μM) do not alter IL-13 induced CREB phosphorylation. N = 3. Scale bar: 20μm. ****: p<0.0001.

Since CaMKII and PKA cascades are strongly activated by glutamate receptors, and glutamate receptors are prominently represented among the targets of IL-13, we hypothesized that CREB phosphorylation may be ultimately dependent on glutamate receptor function. We used two AMPAR, (NBQX and CNQX), an NMADR antagonist (MK-801) and a Voltage-depedent Calcium Channel (VDCC) inhibitor (Mibefradil) to verify our hypothesis. Indeed, IL-13-induced CREB phosphorylation was substantially diminished by AMPAR antagonists NBQX and CNQX as well as by the Voltage-Dependent Calcium Channel blocker Mibefradil but was completely abolished by the NMDAR antagonist MK-801 (Figure 5C-D).

In conclusion, IL-13 activates its cognate receptor IL-13Ra1 at synaptic sites and sets in motion signaling events involving JAK and ERK, as well as NMDA and AMPA glutamate receptors and CaMKII and PKA, ultimately leading to CREB, STAT3 and STAT6 phosphorylation as well as the induction of multiple immediate-early genes.

### 6. IL-13 upregulates synaptic activity

The increases phosphorylation of glutamate receptors and presynaptic proteins, together with the increase in pCREB and the induction of multiple IEGs, provide converging support to the hypothesis that the ultimate effect of IL-13 is to increase neuronal firing. We used the anti-synaptotagmin feeding assay (Catanese et al., 2018) to assess the rate of synaptic activity. In this assay, an antibody against an endoluminal epitope of synaptotagmin was added to the culture medium, so that only the synapses with an active cycling of presynaptic vesicles (only those in which the synaptotagmin luminal epitope is exposed to the external medium before being re-endocytosed) are labelled. Most importantly, IL-13 treatment significantly increased the number of active synaptic terminals after 3h of treatment (Figure 6A-B). Importantly, this effect was blocked by JAK and ERK inhibitors, confirming the selectivity of the readout in relationship with the signaling cascades set in motion by IL-13 (Figure 6A-B).

**Figure 6:**
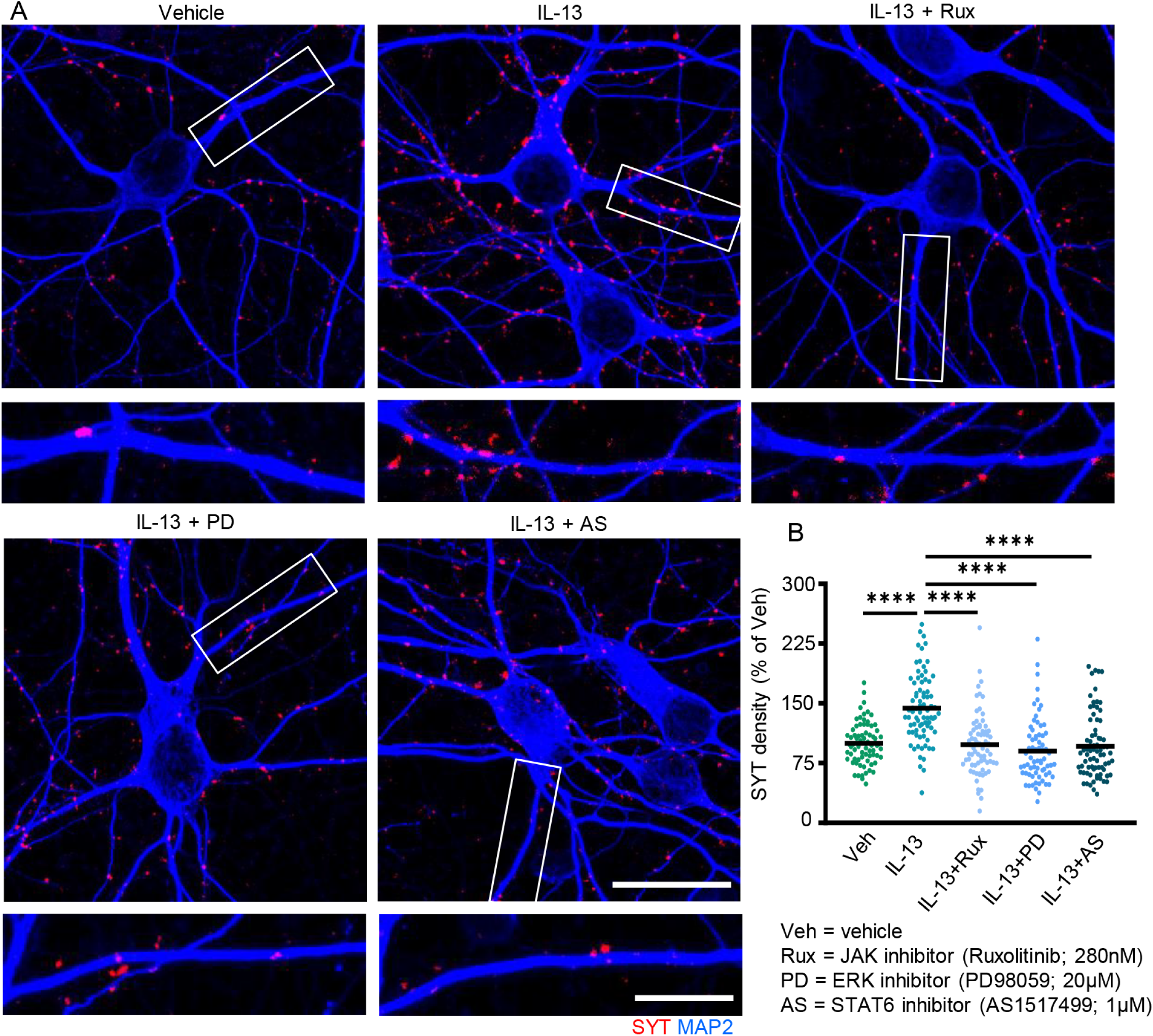
IL-13 increases synaptic activity. A-B. Significant increase of synaptotagmin density after IL-13 treatment (50 ng/ml) in rat cortical neurons. JAK (Ruxolitinib; 280 nM), ERK1/2 (PD98059; 20 μM) and STAT6 (AS1517499; 1 μM) inhibitors significantly abolish the IL-13 induced synaptotagmin density. N = 3. Scale bar overview: 20μm, scale bar insert: 10μm. ****: p<0.0001.

### 7. IL-13 is upregulated upon trauma through activity-dependent and nuclear-calcium-regulated transcription

If IL-13 is a neuron-derived regulator of synaptic function, how is its expression regulated?

In physiological conditions, STAT6 is largely dispensable for IL-13 induction in the brain, since in *STAT6^-/-^* mice the levels of IL-13 protein in the brain are decreased by approximately 17% compared to WT littermates (Supplementary Figure 5B-C).

We explored *in vivo* whether the induction of IL-13 could be dependent on neuronal activity; we used a chemogenetic anion-permeable ion channel (PSAM/PSEM system; Magnus et al., 2011) to inactivate Parvalbumin-positive interneurons, to increase neuronal firing. We performed this manipulation in healthy conditions as well as in the context of acute TBI, since TBI was previously shown to enhance IL-13 expression (olde Heuvel et al., 2019). Injection of the AAV9 in the PV-Cre mice resulted in >95% of PV interneurons expressing the inhibitory PSAM. Silencing PV interneurons (by PSAM expression and PSEM administration) resulted in a significant increase in IL-13 mRNA in neurons compared to control mice (expressing PSAM but administered only the vehicle). Interestingly, TBI alone also upregulated neuronal IL-13 expression, which was not further upregulated by the concomitant chemogenetic silencing of PV interneurons (Figure 7A-B). Thus, increased neuronal activity resulted in enhanced IL-13 transcription at baseline and upon TBI.

**Figure 7:**
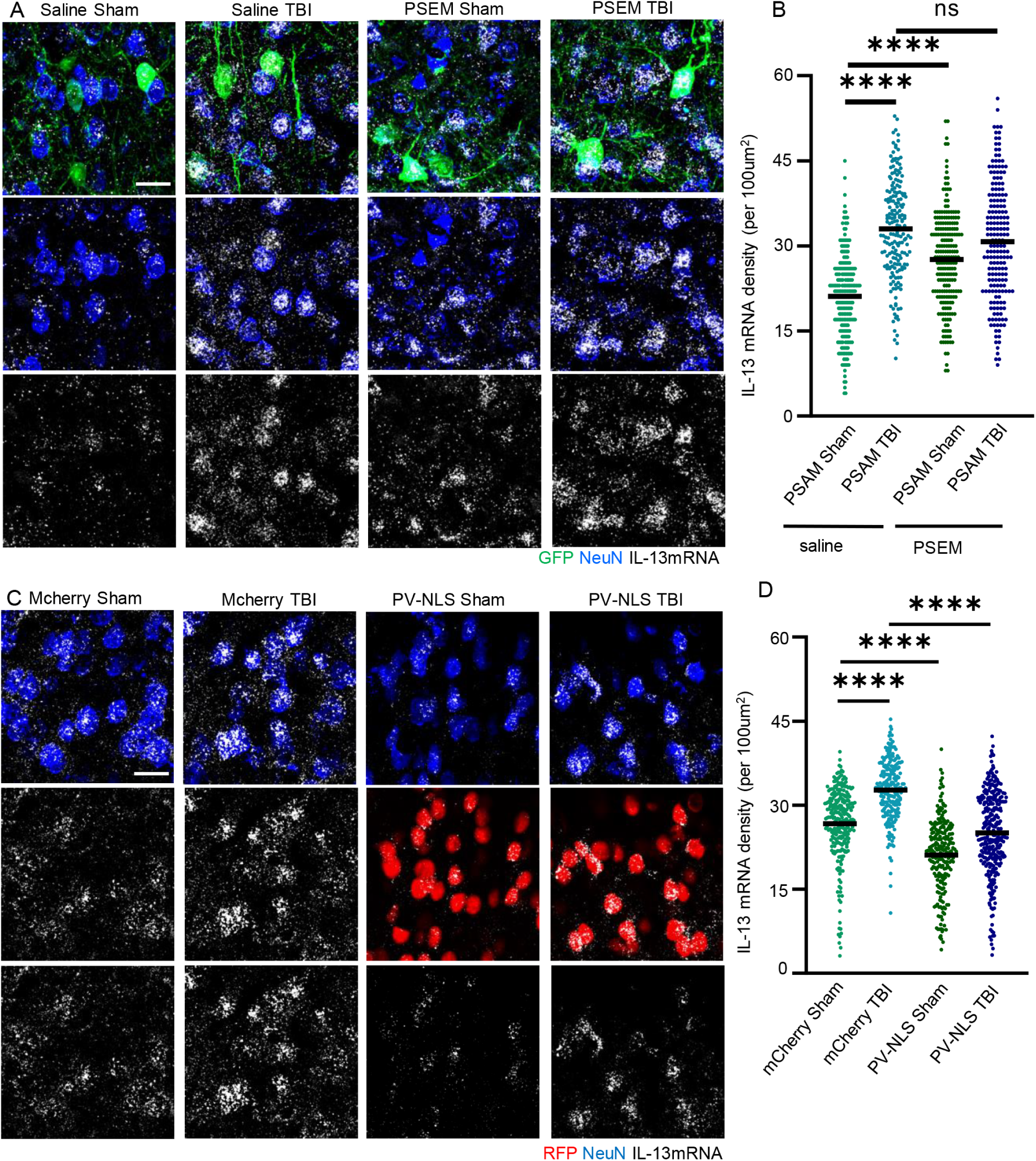
Upregulation of neuronal IL-13 is driven by neuronal activity and nuclear calcium signaling and traumatic brain injury. A-B. Upregulation of IL-13 mRNA expression in a blunt, closed traumatic brain injury murine model (3h timepoint, single-molecule in situ hybridization) Chemogenetic suppression of inhibitory PV interneurons drives the upregulation of IL-13 mRNA at baseline and after trauma. N = 3. Scale bar: 20μm. ****: p<0.0001. C-D. Upregulation of IL-13 mRNA expression in a blunt, closed traumatic brain injury murine model (3h timepoint, single molecule in situ hybridization). Nucleus calcium buffering of principal neurons shows a nuclear calcium dependent transcription of IL-13 at baseline and after trauma. N = 3. Scale bar: 20μm. ****: p<0.0001.

To verify this hypothesis, we set out to block, to a large extent, the nuclear Ca^2+^ signals that are involved in activity-dependent transcription (Bading et al., 2013). We exploited the PV-NLS-mCherry construct (previously described; Mauceri et al., 2015), in which the Ca^2+^- binding protein Parvalbumin is targeted to the nucleus by an attached Nuclear Localization Signal (together with the mCherry tag). The construct was expressed under hSyn promoter and delivered via injection of AAV9 (as control, an empty vector was used).

In sham mice, expression of PV-NLS significantly reduced IL-13 expression compared to the empty-vector. Furthermore, whereas in empty vector-injected mice TBI resulted in the substantial elevation in IL-13 expression, PV-NLS completely blocked this effect. Thus, baseline expression of IL-13 is linked to neuronal activity and nuclear Ca^2+^ signaling, and IL-13 upregulation upon TBI is enhanced by increased neuronal excitability but suppressed by the blunting of nuclear Ca^2+^ signaling (Figure 7C-D).

### 8. IL-13 reduces neuronal sensitivity to excitotoxic cell death

What is the ultimate impact of IL-13 upregulation taking place upon TBI, on neuronal vulnerability? The increase in the phosphorylation of CREB suggests that IL-13 may be beneficial for the cell; conversely, detrimental effects of IL-13 have also been proposed (Mori et al., 2017). We set out to test this hypothesis in an experimentally tractable system. We used a holotomographic microscope (Sandoz et al., 2019) to obtain high-contrast, label-free, time-lapsed imaging of cultured cortical neurons for 12h, acquiring a three-dimensional holographic stack every 15 minutes. Neuronal cultures were exposed to vehicles or to 20µM or 40µM glutamate either alone to simulate an acute excitotoxic environment associated with traumatic injury, or in the presence of IL-13 (50ng/ml; added simultaneously to glutamate). We monitored individual cells for the appearance of nuclear pyknosis (nuclear condensation with substantial increase in refractive index) as a sign of cell sufferance and death; data were recorded for each cell at the timepoint when nuclear pyknosis plateaued. In vehicle-co-treated cultures, exposure to glutamate (20 or 40µM) caused a rapid and relentless increase in nuclear condensation, which plateaued between 3 and 4.5 hours after the exposure; no nuclear pyknosis was observed in vehicle-treated cells (Figure 8A-B and Supplementary Figure 6A-B). Remarkably, co-treatment with IL-13 significantly improved the survival of cultured neurons exposed to either 20 or 40µM glutamate, with median time to piknosis prolonged from 5 to 9.5 hours (Figure 8A-B and Supplementary Figure 6A-B). Remarkably, the beneficial effect of IL-13 was abolished by the co-treatment with inhibitors of IL-13 signaling such as the JAK inhibitor Ruxolitinibas as well as the STAT6 inhibitor AS1517499 (Figure 8A-C).

**Figure 8:**
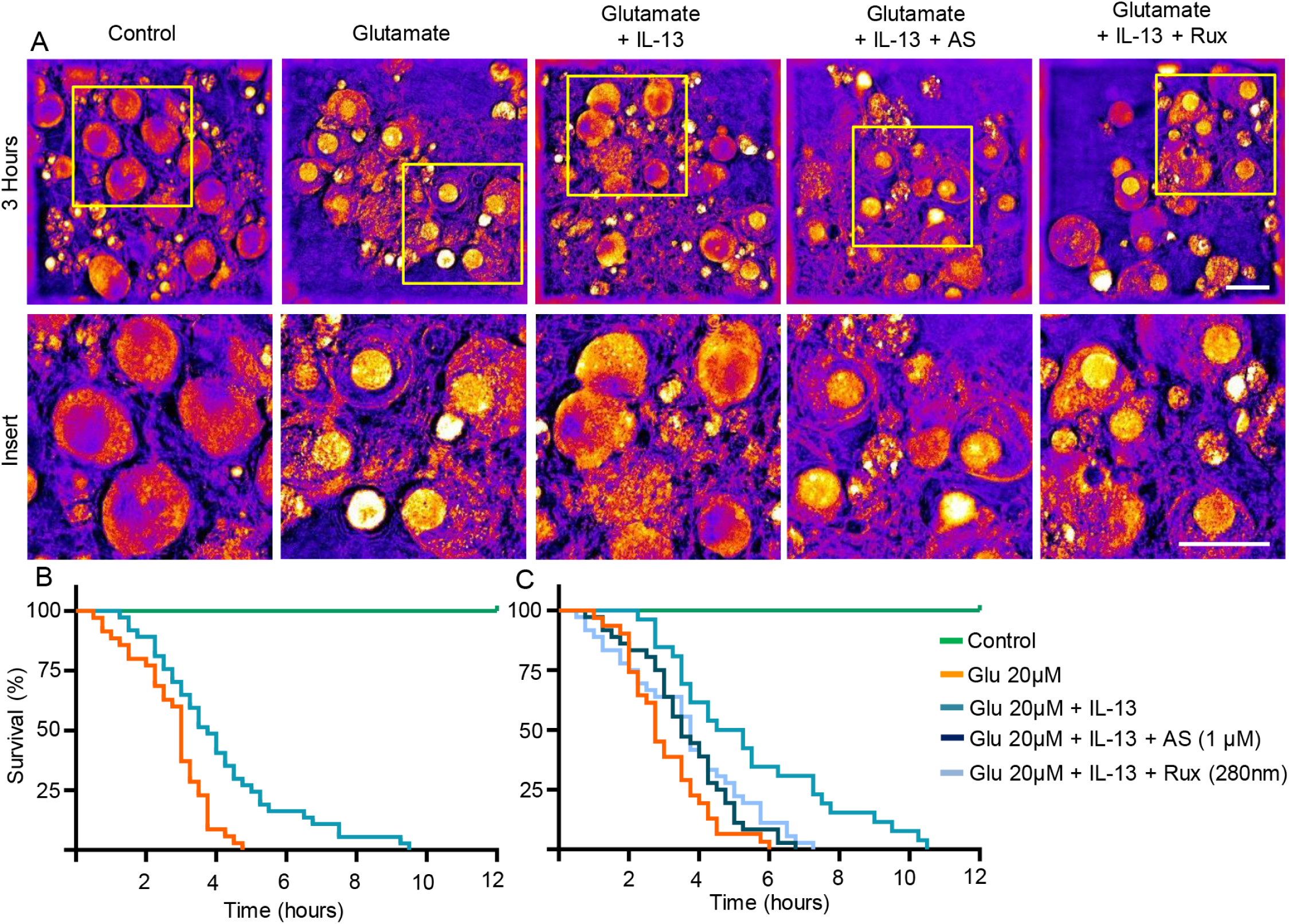
IL-13 reduces excitotoxic neuronal death. A-C. Significant reduction of glutamate (20 μM) induced neuronal toxicity after IL-13 treatment (50 ng/ml) in rat cortical neurons (Holotomographic label-free live cell imaging (Nanolive)). Treatment with JAK (Ruxolitinib; 280 nM) and STAT6 (AS1517499; 1 μM) inhibitors show that the protective effects of IL-13 are dependent on a JAK/STAT6 dependent mechanism. N = 4. Scale bar: 20μm.

### 9. IL-13 and IL-13Ra1 are expressed in human neurons and are upregulated in brain and CSF of TBI patients

Using three distinct approaches, we addressed the relevance of our murine investigations of IL-13 in normal human brain and also in neurotrauma patients.

Firstly, we explored the immunoreactivity for IL-13 and IL-13Ra1 in samples of motor cortex from four independent healthy (i.e., not affected by trauma or neurodegenerative processes, full details are reported in Table 1) post-mortem human brains. IL-13 immunoreactivity was detected in three distinct populations, differing for intensity and location. A large number of moderately positive cells with neuronal morphology was seen across cortical layers, in particular in the upper layers; immunoreactivity was concentrated in the cell body and proximal dendrites whereas a stronger immunoreactivity had characteristics of a punctate staining located along the dendrites and in the neuropil (Figure 9B). A comparatively smaller number of cells with neuronal morphology displayed a strong IL-13 immunoreactivity both in the cell body and in the associated axonal processes, with punctate morphology (Figure 9B). A third population was identified in the form of very strongly IL-13 expressing cells with astroglial morphology located in the white matter underlying the cortical layers. On the other hand, IL-13Ra1 expression appeared more homogeneous in terms of expression intensity, with a large number of cells with neuronal morphology showing immunolocalization of the receptor, with moderate intensity, in the cell body and along the dendrites, up to the apical tuft as well as in basal dendrites (Figure 9C).

**Figure 9:**
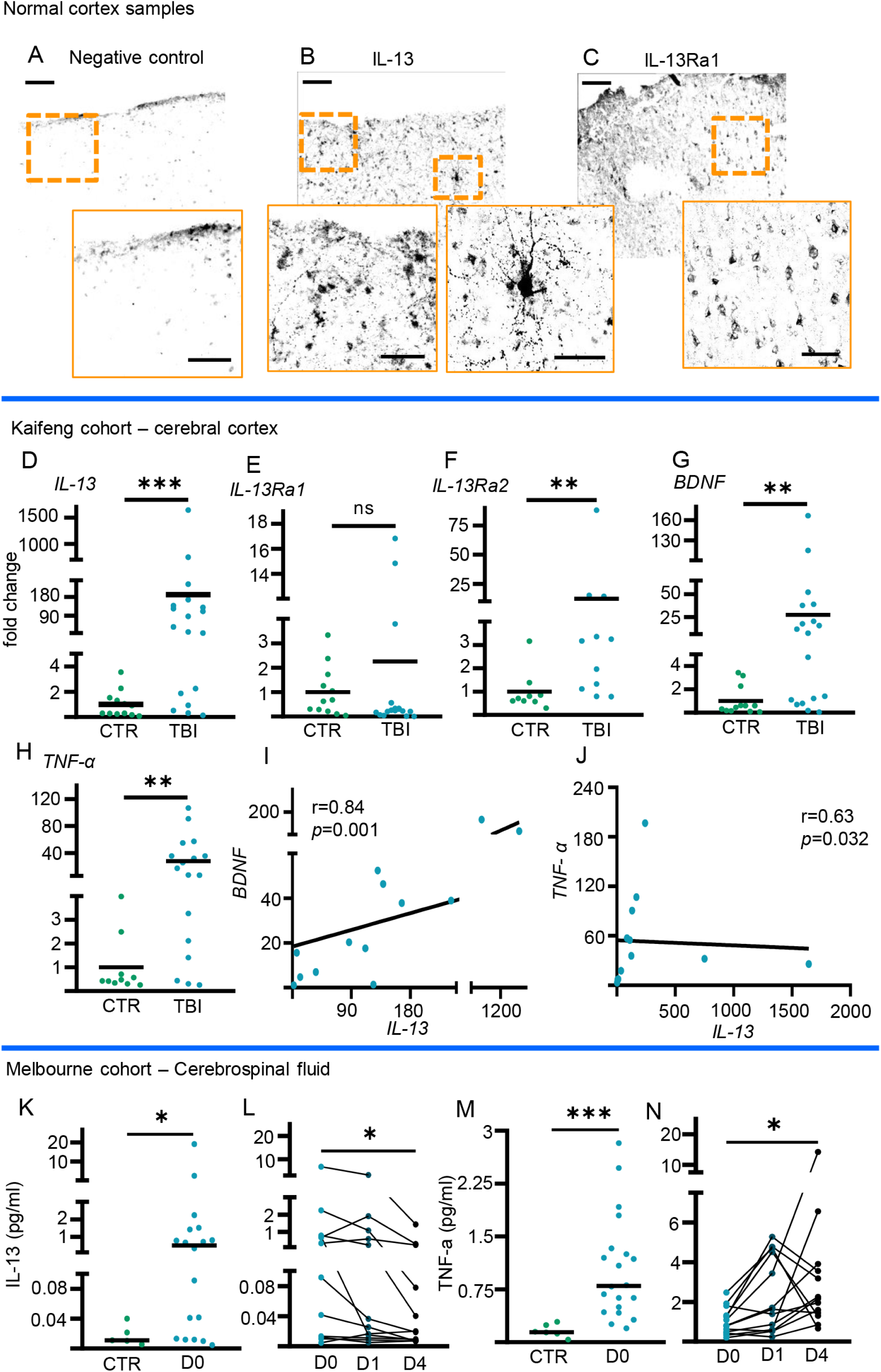
IL-13 is expressed in the human brain and is upregulated upon TBI in the human cortex and CSF. A. Negative-control Immunohistochemical staining (no primary antibody, but secondary added) shows no signal in human post-mortem cortical tissue. N = 3. Scale bar overview: 100μm, scale bar insert: 50μm. B. Immunohistochemical staining of IL-13 shows moderate, high and very high-expressing neuronal populations in human post-mortem cortical tissue. N = 3. Scale bar overview: 100μm, scale bar insert: 50μm. C. Immunohistochemical staining of IL-13Ra1 shows a large number of neurons across all cortical layers of post-mortem cortical tissue. N = 3. Scale bar overview: 100μm, scale bar insert: 50μm. D. Significant upregulation of IL-13 expression in human cortical tissue samples resected after traumatic brain injury (RT-qPCR).. CTR N = 14; TBI N = 19. ***: p<0.001. E. No significant difference of IL-13Ra1 expression in human cortical tissue samples resected after traumatic brain injury (RT-qPCR). CTR N = 14; TBI N = 19. F. Significant upregulation of IL-13Ra2 expression in human cortical tissue samples resected after traumatic brain injury (RT-qPCR). CTR N = 14; TBI N = 19. **: p<0.01. G. Significant upregulation of BDNF expression in human cortical tissue samples resected after traumatic brain injury (RT-qPCR). CTR N = 14; TBI N = 19. **: p<0.01. H. Significant upregulation of TNF-α expression in human cortical tissue samples resected after traumatic brain injury (RT-qPCR). CTR N = 14; TBI N = 19. **: p<0.01. I. Significant correlation between IL-13 and BDNF expression in human cortical tissue samples resected after traumatic brain injury. N = 13. J. Significant inverse correlation between IL-13 and TNF-α expression in human cortical tissue samples resected after traumatic brain injury. N = 13. K. Significant upregulation of IL-13 protein in cerebrospinal fluid samples from traumatic brain injury patients (SIMOA assay) . CTR N = 5; TBI N = 18. *: p<0.05. L. Significant downregulation of IL-13 protein in cerebrospinal fluid samples 4 days after traumatic brain injury compared to the initial increase upon admission (SIMOA assay). D0 N = 12; D1 N = 12; D4 N = 12. *: p<0.05. M. Significant upregulation of TNF-α protein in cerebrospinal fluid samples from traumatic brain injury patients (SIMOA assay). CTR N = 5; TBI N = 20. *: p<0.05. N. Significant upregulation of TNF-α protein in cerebrospinal fluid samples 4 days after traumatic brain injury compared to the initial increase upon admission (SIMOA assay). D0 N = 12; D1 N = 12; D4 N = 12. *: p<0.05.

**Table 1:** Full details normal human cohort (Ulm).

|  |  |
| --- | --- |
| Normal human cohort (Ulm) | Case information |
| N (M/F) | 3 (2/1) |
| Age | 66, 71, 74 |

Altogether our experiments show that the human brain displays a pattern of expression of IL-13 and IL-13Ra1 compatible with that seen in the murine brain, validating our experimental models.

Secondly, we investigated whether the upregulation of IL-13 transcription observed in murine models could be confirmed in human samples. Human cortical samples (n=19) resected in the context during the neurosurgical treatment of acute traumatic injury and, as controls, fragments of healthy human cortex obtained in the context of elective neurosurgery (clipping of unruptured aneurysms, n=14; full details of the two groups -collectively henceforth defined as “Kaifeng Cohort”- are reported in Table 2) were processed to extract the total RNA content for assessment of gene expression. Overall, TBI samples displayed a significantly higher level of IL-13 expression compared to control cortical samples (Figure 9D), in agreement with the observations in the murine model. Interestingly, the expression of IL-13Ra1 was unchanged in TBI samples (Figure 9E), whereas IL-13Ra2 was upregulated upon TBI (Figure 9F). The expression of the inflammatory marker TNF-α and of the neurotrophin BDNF were also upregulated after TBI (Figure 9G-H). A substantial degree of variability was observed among TBI samples, with 5 out of 19 showing very low levels of expression of any of the tested genes, including the housekeeping genes such as gaphd; this was attributed to the possible inclusion of necrotic tissue into the samples area. When these five samples were excluded, we found a significant correlation between IL-13 levels and BDNF expression levels (Figure 9I), whereas no correlation was found with TNF-a (Figure 9J), indicating that elevation of IL-13 was not a consequence of the overall inflammatory response but was correlated with neuronal responses to TBI.

**Table 2:** Full patient details Kaifeng cohort.

| Kaifeng Cohort | **Control group** | **TBI group** |
| --- | --- | --- |
| **N (M/F)** | 14 (9/5) | 19 (13/6) |
| **Age (median, range)** | 73 (42-83) | 57 (23-86) |
| **Glasgow Coma Scale (median, range)** | N/A | 4 (3-8) |
| **Surgery time after TBI (h)** | N/A | 4 (3-6) |

Thirdly, we explored whether the elevation of IL-13 could be replicated in the cerebrospinal fluid (CSF) of moderate-severe TBI patients. We analysed samples obtained from CSF drainage from TBI patients (day0 n=18, day1 n=27 day4 n=28), or, as controls, from patients subjected to CSF drainage for non-traumatic reasons (n=6). This cohort (henceforth collectively defined as “Melbourne Cohort”) is independent of the Kaifeng Cohort and has been the object of publication before (Yan et al., 2014); clinical and demographic characteristics are briefly summarized in Table 3. We used a Single-Molecule Array (SIMOA) platform to determine the concentration of IL-13 and, as proxy of inflammation (Zhou et al., 2021), of TNF-α in CSF samples obtained within 24h of TBI (comparably to the Kaifeng Cohort); for a distinct subset of patients, serial samples were available from day 0 (within 24h of trauma), day 1 and day 4 and these samples were independently analyzed. We found that, compared to controls, TBI patients displayed a very strong elevation, although with substantial variability from case to case, in the level of IL-13 in the acute phase of TBI (<24h; Figure 9K), in agreement with the murine data and the Kaifeng cohort expression data. The analysis of the serial samples showed that in most cases IL-13 levels tended to peak already at day 0 and then progressively declined over time (day 0 vs day 4; Figure 9L). Although TNF-α was also strongly elevated upon TBI (Figure 9M), there was no correlation between IL-13 and TNF-α levels with TNF-α levels peaking (in all but 2 cases) at day 1 or later (Figure 9N).

**Table 3:** Full patient details Melbourne cohort.

| Melbourne cohort | Control group* | D0 group | longitudinal subgroup |
| --- | --- | --- | --- |
| **N (M/F)** | 6 (2/4) | 27 (19/8) | 11 (7/4) |
| **Age (median, range)** | 71 (53-85) | 37 (23-55) | 29 (23-42) |
| **Glasgow Coma Scale (median, range)** | N/A | 6 (3-9) | 6 (3-7) |
| **Injury Severity Score (median, range)** | N/A | 35 (17-59) | 43 (17-59) |
| **Glasgow Outcome Scale-Extended (median, range)** | N/A | 3.5 (1-8) | 3 (1-6) |

Taken together, the findings from the Kaifeng Cohort and the Melbourne Cohort are in remarkable agreement and demonstrate that upon trauma, the cortex and the CSF display a massive but transient increase in IL-13 levels, which are largely uncorrelated with the upregulation of inflammatory cytokines. Therefore, human neurons are exposed to increased levels of IL-13, making our experimental investigation in murine models relevant to human TBI.

## Discussion

In the present study we provide convincing evidence pointing toward a previously unappreciated role of IL-13 as an endogenous regulator of neuronal origin, synaptic biology and neuronal activity. Furthermore, we show that elevation of IL-13 is a common phenomenon taking place in rodents as well as human brains, and that in the condition of brain injury, IL-13 has fundamentally beneficial effects in protecting neuronal survival.

Our data supports the notion that IL-13 neuronal biology shares several similarities with synaptic modulators such as neurotrophins. In fact, IL-13 is upregulated upon increased neuronal firing at baseline and in TBI, and its neural transcription is loosely dependent on STAT6 (in contrast to immune cells) but strongly dependent upon nuclear-calcium signals (Bading, 2013). Furthermore, IL-13 causes a significant increase in NMDA and AMPA receptor phosphorylation, both events associated with the increased recruitment of these glutamate receptors at the synapses and their trafficking to the surface (Roche et al.,1996; Roche et al., 2001; Lussier et al., 2015). In addition, the upregulation of synaptic activity and glutamate receptor activity results in a dramatic elevation of CREB phosphorylation and, in turn, of a number of CREB-regulated genes. Taken together, these data suggest that IL-13 is a previously unknown mediator of synaptic plasticity, displaying activity-dependent potentiation and stabilization. On this ground, it can be hypothesized that loss of IL-13 may produce phenotypes associated with reduced learning and impaired memory retention.

In support of this hypothesis, IL-13^-/-^ mice do display abnormalities often related to disturbances of synaptic plasticity: in fact, IL-13^-/-^ mice performed very poorly in a 4-days Morris Water Maze (MWM) test and almost completely failed at the reversal learning in the same setting (Brombacher et al., 2017). Interestingly, similar, although less prominent, behavioural abnormalities were observed in mice lacking the IL-4 and IL-13 co-receptor (IL-4Ra^-/-^ mice); these mice displayed significantly longer latencies to locate the platform in the MWM test. Although these abnormalities were attributed to the impairment of BDNF secretion by astrocytes under the control of adaptive immunity cells (Brombacher et al., 2017), we propose that failure in synaptic plasticity due to the loss of neuronal IL-13 may substantially contribute to this phenotype. In fact, IL-13 appears to drive phosphorylation events associated with the increased insertion in synapses and increased neuronal activity, both recognised as the hallmarks of the learning process. In this context, additional mechanisms may be involved but remain to be investigated. It could be assumed that IL-13 produced by neurons may reach local microglia through spill-over and may induce a specific microglial phenotype (Littlefield et al., 2017) involved in the remodelling of synaptic contacts (Nguyen et al., 2020).

Our data also shows that neurons exposed to IL-13 display a reduced sensitivity to excitotoxic cell death. At least two mechanisms may be responsible for this effect: firstly, IL-13 could upregulate synaptic signaling leading to CREB phosphorylation, a process well-known to be associated with neuroprotection and reduced neuronal vulnerability (Yan et al., 2020; Hardingham et al., 2002); secondly, after the initial upregulation, IL-13 could lead to a rapid de-phosphorylation of many NMDA and AMPA receptor subunits, conducive to their removal from cell surface and thus effectively blocking glutamate-dependent ion fluxes. Recently, additional mechanisms for beneficial properties of IL-13 in pathological conditions have been proposed, with major focus on the neuroimmunological aspects. In particular, insertion of cells providing a continuous source of IL-13 resulted in a strong M2 polarization of microglia and macrophages upon stroke (Hamzei Taj et al., 2018). A similar shift was observed also upon peripheral administration of IL-13 (Kolosowska et al., 2019). However, it must be noted that in a severe, permanent carotid artery occlusion model of stroke, simultaneous deletion of IL-13, IL-9, IL-4 and IL-5 did not worsen neurological outcome (Perego et al., 2019). In the context of TBI, IL-13 has been shown to attenuate the acute motor deficits in the rotarod test, produce a faster recovery in the foot-fall test and in the wire-hanging tests when administered through intranasal instillation. A decrease in the so-called pro-inflammatory microglial phenotype together with an increased microglial phagocytosis is thought to be involved in these beneficial effects (Miao et al., 2020). The anti-inflammatory properties of IL-13 have been also associated with its ability to trigger the apoptosis of reactive microglia (Yang et al., 2002; Shin et al., 2004). However, in an experimental model of TBI (controlled cortical injury), suppression of IL-13 through the administration of a neutralizing antibody protected neurons from the induction of piroptosis (Gao et al., 2020). Concordant with this detrimental function, it was reported that IL-13 increased oxygen radical production by dopaminergic neurons (Morrison et al., 2012; Maher et al., 2018) and that ablation of neuronal IL-13 signaling (in IL-13Ra1 knock-outs) prevented the loss of dopaminergic neurons in the substantia nigra under chronic stress (Mori et al., 2017). Our findings help to reconcile this conflicting evidence around the role of IL-13: in fact, increased phosphorylation of glutamate receptors and their enhanced membrane localization may amplify the excitatory inputs which, depending on the neuronal type and the concentration, may enhance ROS production and, at the same time, trigger CREB-dependent transcriptional responses. In this regard, here we show that the impact of IL-13 on CREB phosphorylation is not linear but bell-shaped: so that the increase or decrease in CREB activation may depend on the amount of IL-13 available and the resulting cellular response.

In summary, we have used three distinct approaches to demonstrate the relevance of our findings on the role of IL-13 for human TBI. We define IL-13 and IL-13Ra1 immunoreactivity in at least two neuronal populations in the normal human cortex, whose level of expression differed significantly. Since immunostaining in human post-mortem tissue is subject to potential artifacts, it is not possible to conclude that these are indeed distinct cell populations; however, neuronal immunoreactivity for IL-13 and IL-13Ra1 does confirm that murine findings recapitulate at least some aspects of the human biology. The analysis of IL-13 expression in CSF and brain samples from TBI patients further confirms that the upregulation of IL-13 observed in mice does take place in humans as well. Nevertheless, some limitations apply to the use of human tissues. In fact, we found that some samples had small amounts of a variety of mRNAs, suggesting the presence of necrotic tissue. Once those samples were removed, IL-13 levels were highly correlated with BDNF concentrations, whose neuronal induction is dependent on neuronal activity (Rocamora et al., 1996; Bloodgood et al., 2013) In contrast, IL-13 concentrations did not correlate with TNF-α, further indicating that IL-13 upregulation is not a direct consequence of inflammatory responses to TBI. Interestingly, the findings from the TBI patient Kaifeng Cohort were corroborated in the Melbourne cohort, where, once again, IL-13 was found elevated (in agreement with previous reports; Abboud et al., 2016), however without any associations with the levels of TNF-α in CSF. As shown previously for other cytokines measured in the Melbourne cohort (Yan et al., 2014), a significant variability was observed in the concentrations of IL-13 in CSF, which ranged from control levels to almost 100-fold the baseline. Since the Melbourne cohort includes patients with a substantial variability in TBI classification, extent of brain damage and prognosis (Yan et al., 2014), it can be postulated that IL-13 elevation may be characteristic of a subset of patients. To this respect, the role of IL-13 in patient stratification and prognosis is still unresolved and may need a substantially larger cohort size to be established. Nevertheless, the findings of the Kaifeng and Melbourne patient cohorts indicate that human neurons are exposed to high levels of IL-13 upon TBI, both at the site of injury and in the CSF. Therefore, the investigation of the effects of neuronal exposure to IL-13 in in vitro or murine models has translational validity.

The evidence brought forward in this study, identifying IL-13 as a new neuronal regulator of synaptic structure and function, will certainly pave the way to explore a number of unresolved questions around the role of Il-13 in brain physiology and pathophysiology. This is especially relevant in the light of anti-IL-13 therapeutics being developed for allergic conditions as well as against gliomas (Debinski et al., 1999).

Dysregulation of synaptic plasticity and of microglial reactivity are two aspects often encountered in acute and chronic neurological and psychiatric conditions (Bennet and Molofsky, 2019). Our findings reveal that IL-13 may be one part of the machinery involved in both systems and may provide new entry points for therapeutic manipulation of both at once.

## Methods

### 1. Animals

Primary neuronal rat cultures were approved by the Ulm University veterinary and animal experimentation committee under the license number O.103-12. Intracerebral AAV injection, chemogenetics and TBI were approved by the Regierungspräsidium Tübingen under license number 1420.

STAT6^−/−^ mice (B6.129S2(C)-*Stat6* /J) were previously described (Kaplan et al., 1996) and were backcrossed for more than 10 generations to BALB/c background (Symowski and Voehringer., 2019). IL-13^-/-^ mice (Il13t^m1.1Anjm^) were previously reported (McKenzie et al., 1998) and have been bred on a C57/BL6J genetic background for >10 generations (Wodsedalek et al., 2019).

PV-Cre mice (B6.129P2-*Pvalb^tm1(cre)Arbr^*/J), kind gift of Silvia Arber and Pico Caroni, were used for the in vivo TBI experiments and the chemogenetic experiments.

For histology on normal mouse cortex, WT mice (B6SJLF1/J) were used. All mice were group housed from weaning to the time of the procedure, with ad libitum access to food and water, and under a light/dark cycle of 14/ 10 h, at 24 °C and humidity 60–80%.

### 2. Mouse model of traumatic brain injury

Traumatic brain injury was induced using a modified closed, weight drop model (Flierl et al., 2009). Briefly, adult mice aged p60-80 were anesthetized with 5% sevoflurane in 95% O_2_ and treated with 0.1mg/Kg of buprenorphine prior to the injury. The skin was incised on the midline to expose the skull and the mouse was positioned in the weight drop apparatus, in which the head was fixed onto the holding frame. TBI was delivered by dropping a weight of 120 g from a height of 40 cm and a displacement into the skull of 1.5 mm onto the coordinates of the injection site (x = +2.0; y = -2.0; z = 0.0). 100% of 0_2_ was administered after the impact until normal breathing was restored. After the TBI, the skin was sutured with Prolene 6-0 and mice were transferred to their cage with water and food ad-libitum. Mice were euthanized 3 hours post TBI for further experimentation.

### 3. Immunofluorescence staining on mouse brain

Brain samples were processed as previously described (olde Heuvel et al., 2019). Briefly, mice were perfused with 4% PFA in PBS, brain tissue was dissected and post-fixed in 4% PFA overnight. Brain tissue was washed and cryoprotected in 30% sucrose for 2 days, after which the samples were embedded in OCT (tissue tek). 40um sections were cut and subsequently blocked (3% BSA, 0.3% triton-X100, 1x PBS) for 2h at RT, after which primary antibodies (Supplementary Table 1) were diluted in blocking buffer and incubated for 48h at 4□. Sections were washed 3x 30min followed by incubation with secondary antibodies (Supplementary Table 1) diluted in blocking buffer for 2h at RT. The sections were washed 3x 30min and mounted with prolong antifade mounting medium (invitrogen).

### 4. Single-molecule mRNA in situ hybridization

We performed the single-molecule mRNA in situ hybridization as previously reported (Wang et al., 2012) in agreement to the manufacturer’s instructions, (ACDBio, RNAscope, fluorescence *in situ* mRNA hybridization for fixed frozen tissue sections, all reagents/buffers were provided by ACDbio), with minor adjustments (olde Heuvel et al., 2019). In brief, brain sections were affixed on glass slides for histology. Target retrieval was performed by boiling the sections for 5 min in target retrieval solution (provided by ACDbio). Brain sections were washed twice with ddH2O and once with ethanol. After which, the slides were pretreated with protease reagent III at 40°C for 30 min and probe hybridization for IL-13, VGLUT1, VGLUT2 and VGAT was performed at 40°C for 4.5 h. The slides were washed twice with washing buffer for 2 min and then incubated with amplification-1 reagent at 40°C for 30 min. After two washes for 2 min each, slides were incubated with amplification-2 reagent at 40°C for 15 min, followed by a two times washing step. The last amplification was performed with a 30 min incubation with amplification-3 followed by 2 washed for 2 min each. The final detection amplification was performed by incubating the amplification-4 reagent at 40°C for 45 min, and the final washing step was extended to 10 min each time. The co-immunostaining procedure followed immediately after this. The sections were blocked with PBS-/- containing 3% BSA and 0,3% Triton-X 100 at room temperature for 1 h and then incubated with primary antibody (Supplementary Table 1) at 4°C overnight. After being washed with PBS-/- 3x30 min, sections were incubated with secondary antibody (Supplementary Table 1) at room temperature for 2 h, followed by the last round of washing. Finally, the sections were mounted with Fluorogold prolong antifade mounting medium (Invitrogen).

### 5. Dissociated cortical neuronal culture and pharmacological treatments

Each pregnant rat (Sprague-Dawley rats, obtained from Janvier Laboratories) had 6-15 embryos, and both sexes were used. After anesthesia with CO2, adult pregnant rats were sacrificed. Laparotomy was performed to extract the embryos (E17-E18). The heads of embryos were removed into a sterile dish with ice cold Hank’s Balanced Salt Solution (HBSS). After removing the skull. The frontal cortical tissues were manually dissected in ice cold HBSS under stereomicroscopic guidance. The cortical tissues were incubated with 0.25% Trypsin/EDTA (1x) at 37 °C and 5% CO2 for 7-15 min. After three washes with Dulbecco’s Modified Eagle Medium-high glucose (4.5 g/L) (DMEM) with 10% fetal bovine serum (FBS), 1% Glutamine (100x) and 1% Penicillin/Streptomycin (P/S) (DMEM+++), the brain tissues were mechanically dissociated in DMEM+++. The cells were resuspended in DMEM+++ after being filtered by a 100 μm sieve. The cortical cells were plated on ibidi dishes (µ-Dish 35 mm) (for Holotomography Live imaging), 6-wells plate (for cells harvest, WB and qPCR) or coverslips in the 24-wells plate (for Immunofluorescence) with DMEM+++. The dishes, plates and coverslips were coated with Poly-L-lysine (0,05 mg/ml) overnight at 37 °C and 5% CO2. DMEM+++ was replaced with Neurobasal medium supplemented with 2% B-27, 1% Glutamine and 1% P/S (NB+++) after 4-6 h. Cortical cells were grown in a humidified atmosphere at 37 °C containing 5% CO2 and medium was half-renewed weekly. All treatments and experiments were performed on DIV 21. Primary cells were treated with IL-13 at 50 ng/ml for 1 h or 3 h. The inhibitors of transcription factors, ERK inhibitor PD98059 (20 µM, Promega), STAT6 inhibitor AS1517499 (1 µM, Axon Medchem), STAT3 inhibitor Stattic (20 µM, Selleckchem), and JAK inhibitor Ruxolitinib (280 nM, Cayman) were added to the culture medium half an hour before IL-13 exposure. In addition, for pretreatment with the inhibitors of glutamate receptors and calcium channels, primary cultures were exposed to AMPA and kainate receptor blocker CNQX (10 µM, Cayman), AMPA receptor blocker NBQX (10 µM, Cayman), NMDA blocker MK-801 (10 µM, Cayman), and calcium channel blocker Mibefradil (10 µM, Cayman) for half an hour before IL-13 treatment. Similarly, the TrkB receptor antagonist ANA-12 (10 µM, Tocris), PKA inhibitor H89 (10 µM, Tocris), CAMKII inhibitor KN-93 (10 µM, Tocris) Cdk-5 inhibitor (Roscovitine, 10 µM) and GSK-3β inhibitor CHIR 98014 (10 µM, Tocris) were added 30 min prior to IL-13 treatment. A similar volume of only DMSO was added to control wells.

### 6. Immunocytochemistry, confocal imaging and image quantification

Immunofluorescence staining was performed as previously described (Olde Heuvel et al., 2019). Briefly, after the treatment, the coverslips were washed two times with ice cold DPBS-/- and the cells were fixed for 15 min with ice cold 4% PFA added with 0.1 M sucrose. Then cells were washed 2 times with ice cold DPBS-/- again and blocked in PBS-/- with 3% BSA and 0,3% Triton-X 100 at RT for 2 h. Subsequently, cells were incubated with primary antibodies (Supplementary Table 1) diluted in blocking buffer at 4°C for 48 hours. After washing with PBS-/- three times for 30 min, cells were incubated with secondary antibodies (Supplementary Table 1) at RT for 2 h and washed three times for 30 min with PBS-/- at RT again. Finally, the coverslips were mounted on microscope glass slides with Fluorogold prolong antifade mounting medium (invitrogen)

A laser-scanning confocal microscope (Leica DMi8) and fluorescence microscope (Keyence) were used. The Leica microscope equipped with an ACS APO 40x and 63x oil DIC immersion objective was used to acquire images in a 1024x1024 pixel 12-bit format and a Z-stacks of 5 μm (step size of 10 x 0.5 μm). Imaging parameters were set to obtain signals from the stained antibody while avoiding saturation. All fluorescent channels were acquired independently to avoid fluorescence cross-bleed. For each treatment, 6 areas were randomly selected for imaging. Each group was repeated 3-5 times. For c-fos, we used the fluorescence microscope (Keyence) equipped with a 4x and 20x air objective.

Ten optical stacks were imported into ImageJ. The confocal stack (10 optical slices each) collapsed in the maximum intensity projection image. For quantification of receptors on the cell membrane (pIL-13Ra1), the mean gray value of the cell membrane per mature cortical neuron was measured. For transcription factors (pCREB, pSTAT6, pSTAT3, DREAM, ATF-3), the mean gray value of the nucleus per mature cortical neuron was measured. Regarding pERK1/2, the mean gray value of the cytoplasm of each mature cortical neuron was measured. The c-fos positive neurons were manually counted by an operator who turned a blind eye to the treatment group.

### 7. Super-resolution STED imaging and image analysis

Super-resolution STED imaging was performed on a Stedycon module (Abberior instruments, Germany) fixed to a Zeiss microscope fitted with a 100x oil objective. Images were acquired in both confocal and STED mode, imaging parameters were set to obtain signals from the staining while avoiding saturation. All fluorescent channels were acquired independently to avoid cross-bleed. A piece of dendrite was randomly selected and a small stack of 1 micron (0.4 micron stack size) was acquired. Images were analysed with the ImageJ software by plotting intensity profiles over two adjacent or overlapping spots with the dendrite as starting point. The distance from post- and presynaptic markers (PSD-95 and Bassoon respectively) was measured.

### 8. Brain fractionation and Western blot

Brain fractionation was acquired as described previously (Reim D et al., 2017). After dissection, the cortical tissue was suspended in Buffer 1 and homogenized with the Teflon douncer. 75 µl sample (Ho) was taken into a new tube called Ho. To remove extracellular matrix, nuclei and cell debris, the obtained homogenate (Ho) was centrifuged at 4°C at 500 x g for 5 min. The supernatant (S1) was collected in a new tube called P3. A 75 µl sample (S1) was taken into a new tube called S1. P3 was centrifuged at 4°C at 10000 x g for 15 min to separate the crude membrane fraction (P2) from the cytosol (S2). The supernatant (S2) was collected. The pellet (P2) was resuspended in 500 µl Buffer 2. P2 was centrifuged at 4°C at 20000 x g for 80 min. The supernatant (S3) contained the soluble synaptic cytosol. The pellet (P3) containing the insoluble PSD fraction was resuspended in Buffer 3.

BCA Protein Assay Kit was used to determine the protein concentrations as described in the manufacturer’s instructions. 40-50 µg protein with 1 x loading buffer and 5% 2-Mercaptoethanol was heated at 95°C for 5 min. Samples were loaded on 8-12% SDS-PAGE gel and transferred to the nitrocellulose membrane. The membranes were blocked and then incubated with primary antibodies (Supplementary Table 1) on the shaker at 4°C overnight. After washing 5 times for 5 min with TBST on the shaker at RT, the membranes were incubated in horseradish secondary antibodies (Supplementary Table 1) in blocking buffer on the shaker at RT. The membranes were washed 5 times for 5 min again. Proteins bound to western blotting membranes were detected by Enhanced chemiluminescence (ECL). ImageJ was used to acquire the mean gray value of the bands.

### 9. RNA extraction and qPCR

RNA isolation was performed as described in the manufacturer’s instructions (QIAzol, Qiagen). The dishes were washed three times with ice cold DPBS-/-. 0.5 ml QIAzol Lysis Reagent was added per well in the 6-well plate. The pipette was used to rinse the bottom of dishes until the cells lysate was uniformly homogeneous. The tube containing the homogenate was placed at RT for 5 min. 0.15 ml chloroform was added and the tube was shanked vigorously for 15 sec. The tube containing the homogenate was kept at RT for 10 min. After centrifuged at 4°C at 12,000 x g for 15 min, the upper, aqueous phase was transferred into a new tube. The RNA was precipitated by mixing with 250ul isopropanol by vertexing. After being kept at RT for 10 min, the tube was centrifuged at 4°C at 12,000 x g for 10 min. The supernatant was discarded. After adding 1 ml 75% ethanol, the tube was centrifuge at 4°C at 8000 x g for 10 min. The supernatant was removed completely, and the RNA pellet was air-dry until the pellet looked like the colour of wet sugar. The RNA pellet was dissolved in 15-25 μl of RNase-free water. The concentration and quality were detected by NanoDrop 2000. The ratio of A260/A280 should be greater than 1.8. Reverse transcription was performed as described previously (Olde Heuvel et al., 2019). 1 μg RNA was mixed with RNase-free water and 5 μl Random primers, dn6, to bring the total volume to 40 μl. The samples were incubated for 10 minutes at 70°C. 15 μl master mix was added, containing 12 μl RT 5x buffer, 2 μl dNTPs, 0.5 μl RiboLock Rnase Inhibitor, 0.5 μl M-MLV RT RNase. After incubating for 10 minutes at RT, the samples were incubated at the water bath at 42°C for 45min. To stop the reaction, the samples were heated at 99°C for 5 min. The cDNA was used immediately for qPCR or store at -20 °C. We used the Light Cycler 480II in this project. 2 μl cDNA was mixed with 5 μl TB Green Premix Ex Taq and 3 μl primer mix (Supplementary Table 2) in a 96-well plate. All the samples were duplicated 6 times. We used the housekeeping gene GAPDH as a control. The CT values obtained from the light cycler were calculated with the 2^-ΔΔCt^ formula.

### 10. Antibody phospho-array assay

Neuroscience Phospho Antibody Array was performed as described in the manufacturer’s instructions (Full Moon BioSystems). After three times washing with ice-cold DPBS-/-, 50 ul/6-well lysis buffer (RIPA + Protease inhibitor 3x + PhosSTOP3x) was added and the cells were scraped on ice. The samples were treated with sonication on ice (5 sec, 5 x 10% cycle, 65% power). The tube with samples was kept on ice for 30 min. After being centrifuged at 4°C at 12,000 x g for 20 min, the supernatant was gently transferred into a new tube. The concentration of protein was detected by BCA Protein Assay Kit as well. 100 µg of protein was used for this array. The volume of the sample was brought to 75 ul by Labeling Buffer. After adding 3 µl of the Biotin/DMF solution, the mixture was incubated at RT for 2 h with vortexing every 10 min. 35 µl of Stop Reagent was added to stop labeling. The Antibody Arrays were blocked with blocking solution for 45 min on the shaker rotating at 55 rpm at RT. After blocking, the slide was washed with Milli-Q grade water ten times. The biotinylated sample was combined with 6 ml Coupling Solution. The slide was incubated with protein coupling mix on the orbital shaker rotating at 35 rpm for 2 h at RT. After coupling, the slide was washed with 1 x Washing Solution on the shaker rotating at 55 rpm for 3 x 10 min. The slide was extensively rinsed with Milli-Q water as before. The slide was submerged by Detection Buffer with 0.5 mg/ml Cy5-streptavidin on the shaker rotating at 35 rpm for 20 min at RT in the dark. The slide was washed by Washing Solution and rinsed with Milli-Q water as before. After being dried by nitrogen, the slide was scanned using a GenePix 4000B array scanner (Molecular Devices, LLC).

For array data analysis, the raw intensity values for each phosphorylated epitope were recorded automatedly via Image recorder software or manually using ImageJ software. The raw data files were loaded in R software and the dataset for each array was preliminary subjected to quality control assessment (QCA); outlier identification, data distribution, intra-array and inter-array normalization. Modified linear modeling-based analysis was then applied to the data to identify targets showing a significant increase or decrease in phosphorylation at the different timepoints. For protein array analyses, the code has been made publicly available on open-access GitHub repository PROTEAS (PROTein array Expression AnalysiS; <u>github.com/Rida-Rehman/PROTEAS</u>).

### 11. Anti-synaptotagmin-1 antibody feeding assay

We performed the anti-synaptotagmin antibody feeding protocol, as previously reported, to detect the synaptic vesicle release rate (Catanese et al., 2018). Briefly, a monoclonal antibody (diluted 1:500) directed against a luminal epitope of Synaptotagmin-1 fluorescently labeled with Oyster®550 (105 103C3, Synaptic System) were added to the culture medium 30 min before fixing. For IL-13 and inhibitors exposure experiments, the same protocol as reported above was applied. After being washed twice with DPBS-/-, the cells were fixed with 4% paraformaldehyde containing 4% sucrose for 10 min. Then the processed primary cells were used for immunofluorescence steps.

### 12. AAV Vectors and Chemogenetics

AAV9 mediating the expression of Pharmacologically Selective Activation Module (PSAM) (Magnus et al., 2011; Saxena et al., 2013) were obtained from Vector Biolabs (Malvern-PA, US), at the titers of 6 × 10^12^ viral genomes/ml, encoding the pAAV-cba-flox-PSAM (Leu141Phe, Tyr116Phe) GlyR-WPRE, anion-permeable (inhibitory PSAM, henceforth inhPSAM) channels. PSEM308, the PSAM agonist, which was obtained from Apex Scientific Inc. (Stony Brook-NY, USA) and dissolved in sterile saline, was administered by intraperitoneal injection 30 min before TBI, at the dose of 5 μg/g.

pAAV-hSyn-PV-nuclear localisation sequence (NLS)-mC, which was constructed by PCR-amplifying the PV-NLS-mCherry coding sequence from pAAV-CMV-PV-NLS-mCherry and then subcloning it into a pAAV-hSyn express, drove the expression of PV-NLS-mCherry under control of the human synapsin promoter, ion plasmid (Schlumm et al. 2013).

### 13. Intracerebral Injection of Viruses

Intracortical injection of AAVs was performed, as previously reported (Chandrasekar et al., 2019), in mice aged P30–P35. Mice were administered buprenorphine (0.05 mg/kg; Reckitt Beckshire Healthcare, Beckshire, UK) and meloxicam (1.0 mg/kg; Böhringer Ingelheim, Biberach an der Riß, Germany) 20 min before the surgery. Mice were positioned into a stereotactic frame under continuous isoflurane anesthesia (4% isoflurane in 96% O2). After incising the scalp at the midline, a burr hole was drilled at the coordinates x = +2.0, y = − 2.0 by a hand micro-drill, which corresponded to the somatosensory cortex. Approximately 200– 500 nL of the viral suspension which was mixed with an equal volume of 1.5% Fast green solution, was injected by a pulled glass capillary which connected to a Picospritzer microfluidic device, over a span of 10 min. In case of preventing the backflow of the virus, the capillary was kept in place for another 10 min. To allow the bone to heal, the burr hole was left open. Prolene 7.0 surgical thread was used to suture the skin. Animals were transferred to a recovery cage with a warmed surface and ad libitum access to food and water after surgery. During the following 72 h, animals were administered additional doses of buprenorphine. The animals were monitored for possible neurological adverse events.

### 14. Holotomography Live imaging

The holotomographic microscope (Nanolive) was used to obtain high-contrast, label-free, time-lapsed imaging of cultured cortical neurons for 12 h, acquiring a three-dimensional holotomographic stack every 15 min. During the imaging, neuronal cultures (grown on optical-grade 35-mm dishes; Ibidi, Munich) were kept at 37°C in a 95/5% O_2_/CO_2_ environment. Neuronal cultures were exposed to vehicles or, to simulate an acute excitotoxic environment associated with traumatic injury, to 20 µM or 40 µM glutamate (added to the culture no more than 5 min before the beginning of the time-lapse imaging) either alone or in presence of IL-13 (50 ng/ml; added at the same time of glutamate addition). We treated the cells with additional STAT6 inhibitor AS1517499 (1 µM, Axon Medchem) and JAK inhibitor Ruxolitinib (280 nM, Cayman) 30 min before the glutamate treatment.

### 15. Human patients’ cohort

The recruitment of patients and the use of brain and CSF samples were authorized by the Kaifeng Central Hospital and Alfred Hospital Human Ethics Committee with approval No. 194-05 and the analysis at Ulm University was authorized by the Ulm University Ethical committee. Normal human cortex samples were obtained in agreement with the procedures approved by the Ulm University ethical committee with approval No. 245/17 (Full details available in Table 1)

P of Kaifeng Cohort were recruited at the Kaifeng Central Hospital, China, between January 2018 and January 2021; informed consent was obtained from the next of the kin. Inclusion criteria were severe TBI with a post-resuscitation GCS ≤8 and the need for neurosurgery to remove hematoma or injured brain tissue. In the TBI cohort, brain samples were collected in the proximity of, but not part of the macroscopic necrotic or hemorrhagic lesion (as visually identified by the neurosurgeon) within 6.5 h after TBI and kept at -80°C. Exclusion criteria comprised pregnancy, neurodegenerative diseases, HIV and other chronic infection/inflammatory diseases, or history of TBI. Inclusion criteria for controls were: non brain-injury or other insults such as cerebral hemorrhage, aneurysm or autoimmune, inflammatory, infectious or neurodegenerative diseases (Full details available in Table 2).

Patients for Melbourne Cohort were recruited at the Alfred Hospital, Melbourne; informed consent was obtained from the next of the kin. Inclusion criteria were: severe TBI with a post-resuscitation GCS ≤8 (unless initial GCS>9 was followed by deterioration requiring intubation) and, upon CT imaging, the need for an extraventricular drain (EVD) for ICP monitoring and therapeutic drainage of CSF. CSF was collected over 24 h and kept at 4°C; samples were obtained on admission (day 0) and daily up to day 5 after injury. Within an hour from collection, samples were centrifuged at 2000g for 15 min at 4°C and stored at - 80°C until analysis. Exclusion criteria comprised pregnancy, neurodegenerative diseases, HIV and other chronic infection/inflammatory diseases, or history of TBI. Out of the 42 TBI patients constituting the original cohort (Yan et al., 2014), we selected samples from 30 patients, depending on the availability of three aliquots for day 0, day 1 and day 4 after injury (Full details available in Table 3).

### 16. Single-molecule Array determination of cytokines in CSF

Single-molecule arrays were performed as described as per manufacturer instructions (Quanterix) for the quantification of TNF-α and IL-13. CSF samples were diluted 1:5 for TNF-α measurements and undiluted for IL-13 measurements.

### 17. Immunohistochemistry of human cortex and imaging

Immunohistochemistry was performed on 50 µm thick cortical sections as described before (Braak et al., 2018). Briefly, sections underwent an antigen retrieval step in citrate buffer (pH=6) for 20 min at 80°C, followed by a blocking step in 5% BSA and 0,25% Triton-X 100 at RT for 90 min. After blocking, sections were incubated in primary antibody (Supplementary Table 1) overnight at RT diluted in Tris buffer. The following day, the sections were washed 3 times with Tris buffer and incubated at RT for 90 min with the corresponding secondary biotinylated antibody diluted in Tris buffer (Supplementary Table 1). The sections were then washed 3 times and treated with avidin-biotin complex solution (VECTASTAIN® Elite ABC-HRP Kit, Vector Lab, #PK-6100) for 90 min at RT. The color reaction was then conducted using 3,3-diaminobenzidine (DAB) solution. Sections were dehydrated gradually in ethanol series, cleared in Xylene twice, and finally mounted on microscope glass slides with Histomount mounting medium (National Diagnostics). The sections were imaged using a brightfield microscope (Keyence, BZ-X800), equipped with a 100x oil objective, by making a tile scan image of the grey and white matter of the cortex.

### 18. Statistical Analysis

Data analysis was performed using Graphpad prism version 8 software. All data was tested for normality using the Shapiro-Wilks test. Grouped analysis was performed using ANOVA with Tukey multiple correction or Kruskal-Wallis test with Dunn’s multiple correction, depending on normality. Kaplan-Meier curves were analysed using the Log-rank (Mantel.Cox) test. When appropriate an unpaired t-test or Mann-Whitney test was used for comparing two groups. Correlation analysis was done by Pearson r or Spearman r, depending on normality. Timeline in the human ‘Melbourne cohort’ was analysed using the Friedman test with Dunn’s multiple correction. Data was shown as mean ± SD and statistical significance was set at p < 0.05.

## Supporting information

supplementary statistics file

## Conflict of interest

The Authors declare no conflict of interest.

## Acknowledgements

This work has been supported by the Deutsche Forschungsgemeinschaft as part of the Collaborative Research Center 1149 “Danger Response, Disturbance Factors and Regenerative Potential after Acute Trauma” (DFG No. 251293561). FR is also supported by DFG through the individual grants no. 443642953, no. 431995586 and no. 446067541, and by the ERANET-NEURON initiative “External Insults to the Nervous System” as part of the MICRONET consortium (funded by BMBF: FKZ 01EW1705A). SL is supported by the China Scholarship Council (Personal certificate No. 201908320357). We are grateful to prof. Stefan Just and to prof. Anita Ignatius for the use of the confocal microscope and of the histology laboratory. We would like to thank the colleagues of the German Center for Neurodegenerative Disease (DZNE)-Ulm for advice and discussion. Technical support by Thomas Lenk and Gizem Yartas was highly appreciated. We would like to acknowledge Ms Ann Sutherland from Victorian State Trauma Outcomes Registry and Department of Epidemiology, Monash University, and Ms Shirley Vallance and Ms Lynne Murray from the Intensive Care unit, Alfred Health for providing the GOSE scores.

## Authors contribution

FR conceived and designed this study. SL and FoH performed all the in vitro experiments, and SL, AF and FoH performed the in vivo TBI experiments. RR performed the bioinformatic analysis of the arrays. OA and SW performed the immunohistochemical study of the human brains. ZL and WZ contributed the data from the Kaifeng cohort brain samples. AC and CMK contributed the CSF samples data from the Melbourne cohort. BK contributed the mRNA expression data from in vitro experiments. CoM, DV and RLR contributed data from transgenic mice. JK reviewed the clinical data from the Kaifeng and the Melbourne cohorts. SL, FR, AL, MHL, BK, CMK, TB and FR wrote the initial draft of the manuscript. FR, TB, AL, MHL, JK and CMK contributed to the final version of the manuscript.

## Figure legends

**Supplementary Figure 1: Verification of neuronal IL-13 and IL-13Ra1 expression and synaptic localisation.**

A. IL-13 mRNA expression in VGLUT1 positive glutamatergic neurons in layer IV of mouse cortical sections (single molecule in situ hybridisation). N = 3. Scale bar: 20μm.

B. increased IL-13 mRNA intensity in VGLUT1 positive glutamatergic neuronal population compared to the global cell population layer IV of mouse cortical sections. N = 3.

C. IL-13 mRNA expression in VGLUT2 positive glutamatergic neurons in layer IV of mouse cortical sections (single molecule in situ hybridisation). N = 3. Scale bar: 20μm.

D. increased IL-13 mRNA intensity in VGLUT2 positive glutamatergic neuronal population compared to the global cell population layer IV of mouse cortical sections. N = 3.

E. Synaptic localisation of another IL-13 antibody in mouse cortical sections (Immunostaining with MAP2, pan-VGLUT and SHANK2). Scale bar overview 10μm and insert 5μm.

F. Baseline mRNA levels of IL-13, IL-13Ra1 and IL-13Ra2 in rat primary cortical neurons (RT-qPCR). N = 3.

G. STED images of single synapses show multiple IL-13Ra1 puncta in a single PSD-95 spot. Scale bar overview: 1μm, scale bar insert: 500nm.

H. STED images of single synapses show multiple IL-13 puncta in a single Bassoon spot. Scale bar overview: 1μm, scale bar insert: 500nm.

**Supplementary Figure 2: Validation of IL-13 antibody and IL-13 probe used for IF and ISH.**

A-B. IL-13 antibody validated on IL-13-/- and IL-13+/+ littermates together with MAP2 and SHANK2 immunostaining. N = 3. Scale bar: 10μm.

C-D. IL-13 ISH probe validated on IL-13-/- and IL-13+/+ littermates together with VGLUT1 probe and counterstained with DAPI. N = 3. Scale bar: 20μm.

**Supplementary Figure 3: IL-13 induces STAT phosphorylation and immediate-early genes transcription.**

A-B. Significant up-phosphorylation of STAT6 in rat cortical neurons 1h and 3h after IL-13 treatment (50 ng/ml) (immunostaining). N = 3. Scale bar: 20μm. ****: p<0.0001.

C-D. Significant up-phosphorylation of STAT3 in rat cortical neurons 1h after IL-13 treatment (50 ng/ml) (immunostaining). N = 3. Scale bar: 20μm. ****: p<0.0001.

E-F. Significant up-phosphorylation of ATF-3 in rat cortical neurons 1h and 3h after IL-13 treatment (50 ng/ml) (immunostaining). n = 3. Scale bar: 20μm. ****: p<0.0001.

G. Significant increase of c-fos, fosB, egr1, gadd45a and gadd45b expression 1h after IL-13 treatment (50ng/ml), with a subsequent decrease 3h after treatment (RT-qPCR). N = 6. *: p<0.05. **: p<0.01. ***: p<0.001.

**Supplementary Figure 4: IL-13Ra1 phosphorylation occurs mainly at synaptic sites.**

A-D. Significant up-phosphorylation of IL-13Ra1 at mature synapses 3h after IL-13 treatment (50 ng/ml) (Immunostaining with MAP2, pan-VGLUT and PSD-95). N = 3. Scale bar overview 10μm and insert 5μm. *: p<0.05. **: p<0.01. ***: p<0.001. ****: p<0.0001.

**Supplementary Figure 5: STAT6 transcriptional response mediates IL-13 expression.**

A. Upregulation of IL-13 mRNA expression in a blunt, closed traumatic brain injury murine model (3h timepoint, single molecule in situ hybridization). N = 3. Scale bar: 50μm.

B. Significant downregulation of IL-13 protein in STAT6-/- mice, IL-13Ra1 is not affected in STAT6-/- mice (Western Blot). N = 3. *: p<0.05.

**Supplementary Figure 6: IL-13 reduces high dose excitotoxic neuronal death.**

A-B. Significant reduction of glutamate (40 μM) induced neuronal toxicity after IL-13 treatment (50 ng/ml) in rat cortical neurons (Holotomographic label-free live cell imaging (Nanolive)). N = 4. Scale bar: 20μm.

**Supplementary Table 1: List of antibodies and other reagents used for immunostaining (IF), immunohistochemistry (IHC), Fluorescent in Situ Hybridisation (RNAscope) or for western blot (WB).**

**Supplementary table 2: List of genes and sequences used for RT-qPCR.**

## Notes

### Competing Interest Statement

The authors have declared no competing interest.

