## Supplementary figures and images for "Interleukin-13 and its receptor are synaptic proteins involved in plasticity and neuroprotection"

### supplementary figure 1

Supplementary_Figure_1.pdf
1562 x 2530

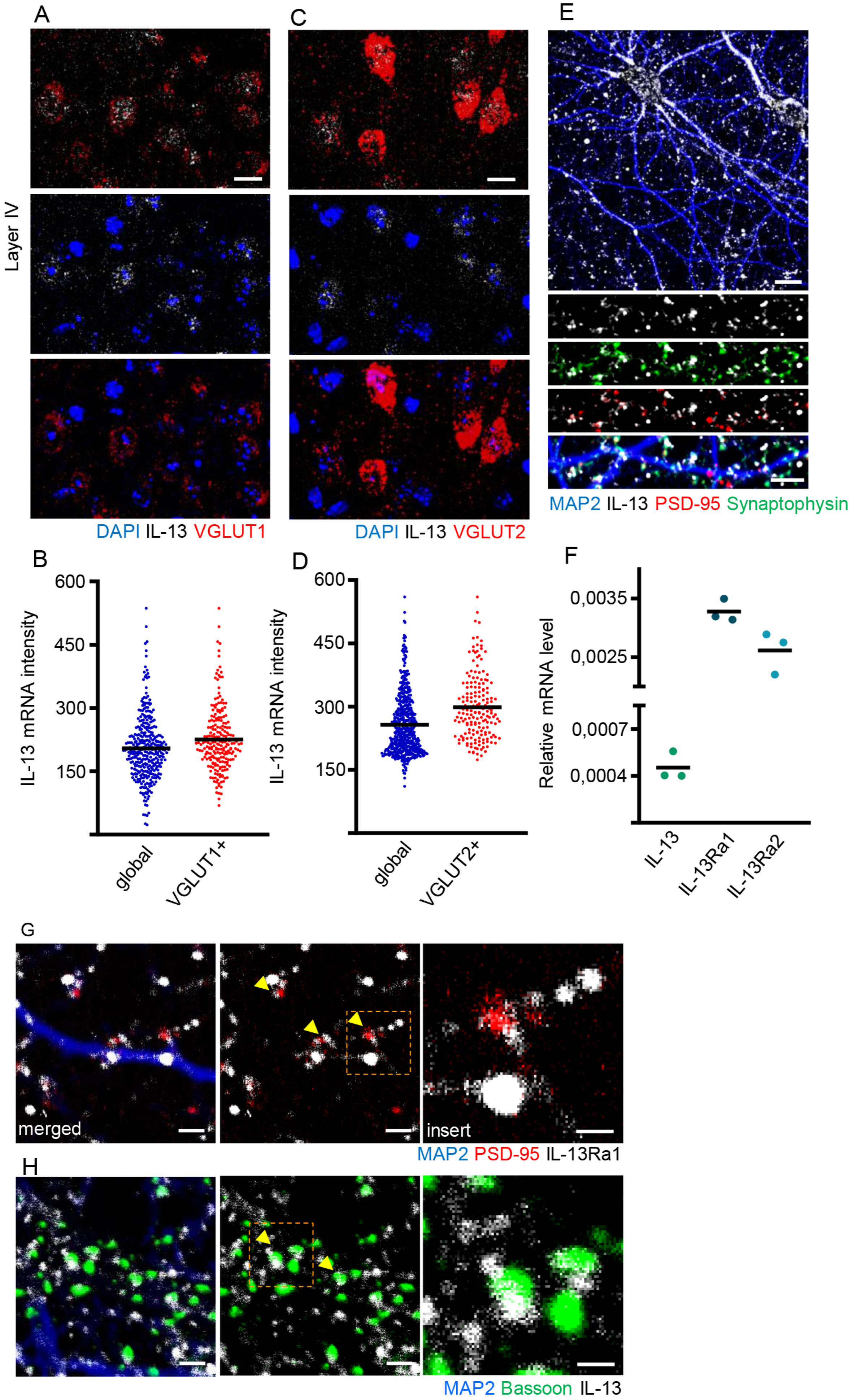

### supplementary figure 2

Supplementary_Figure_2.pdf
1418 x 938

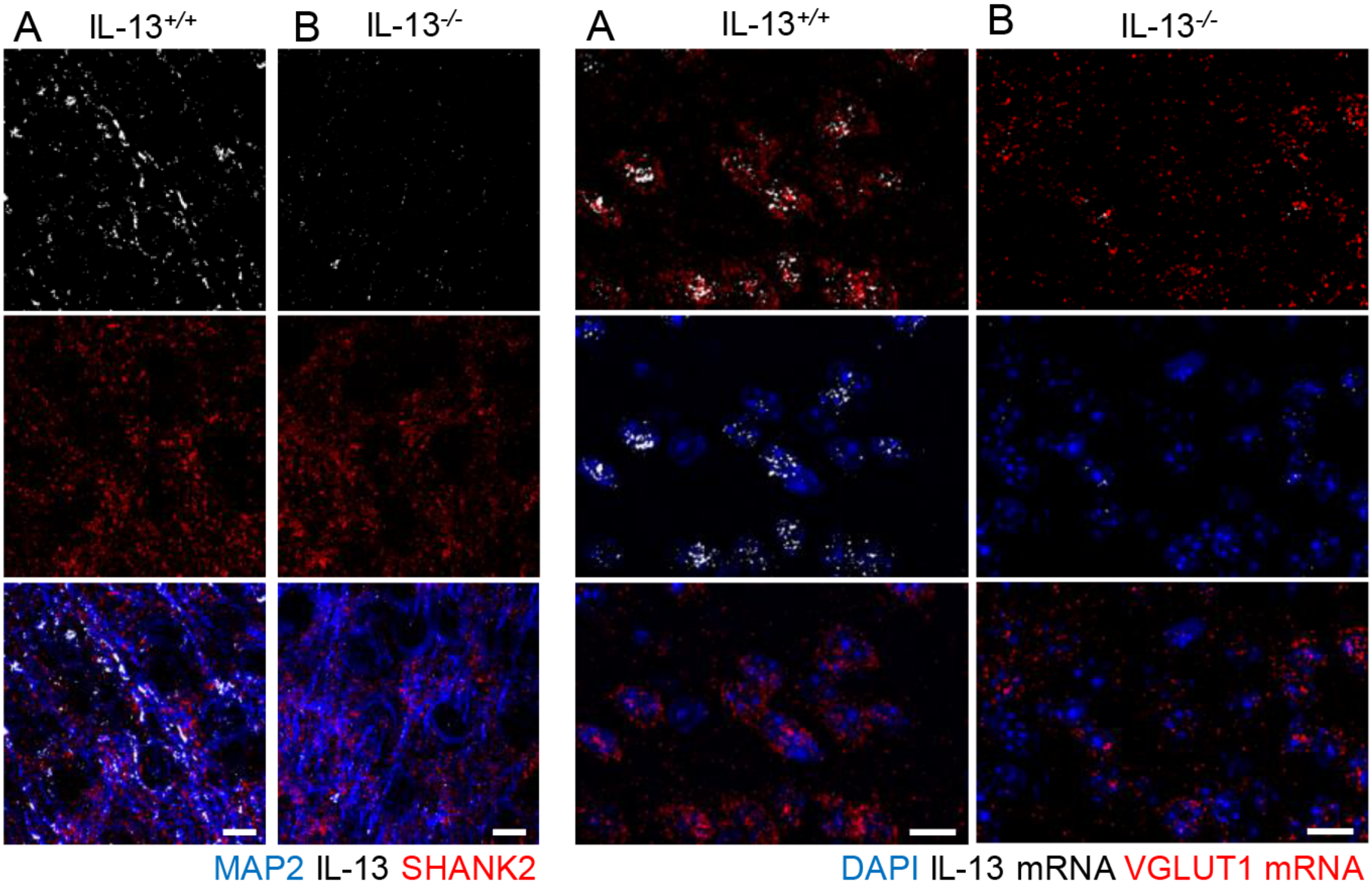

### supplementary figure 3

Supplementary_Figure_3.pdf
1045 x 2298

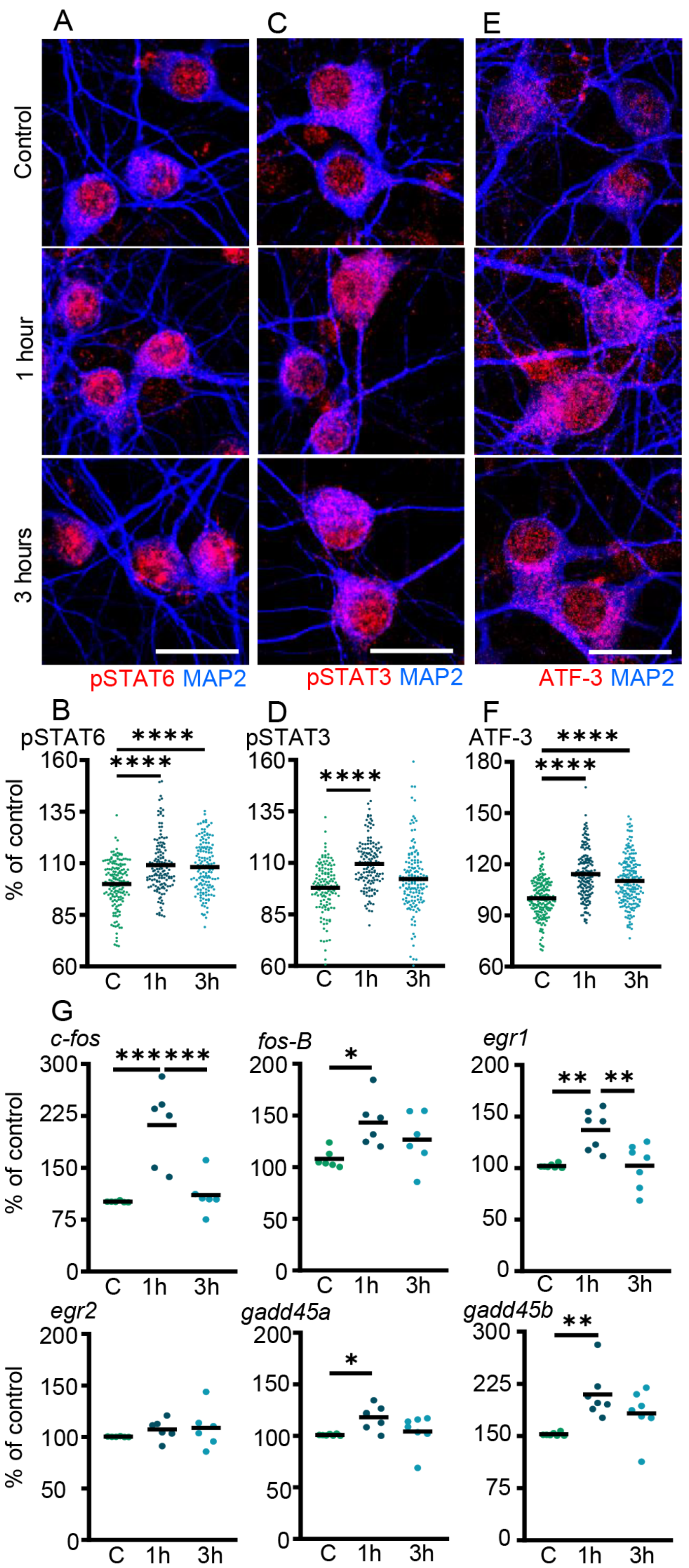

### supplementary figure 4

Supplementary_Figure_4.pdf
1170 x 1242

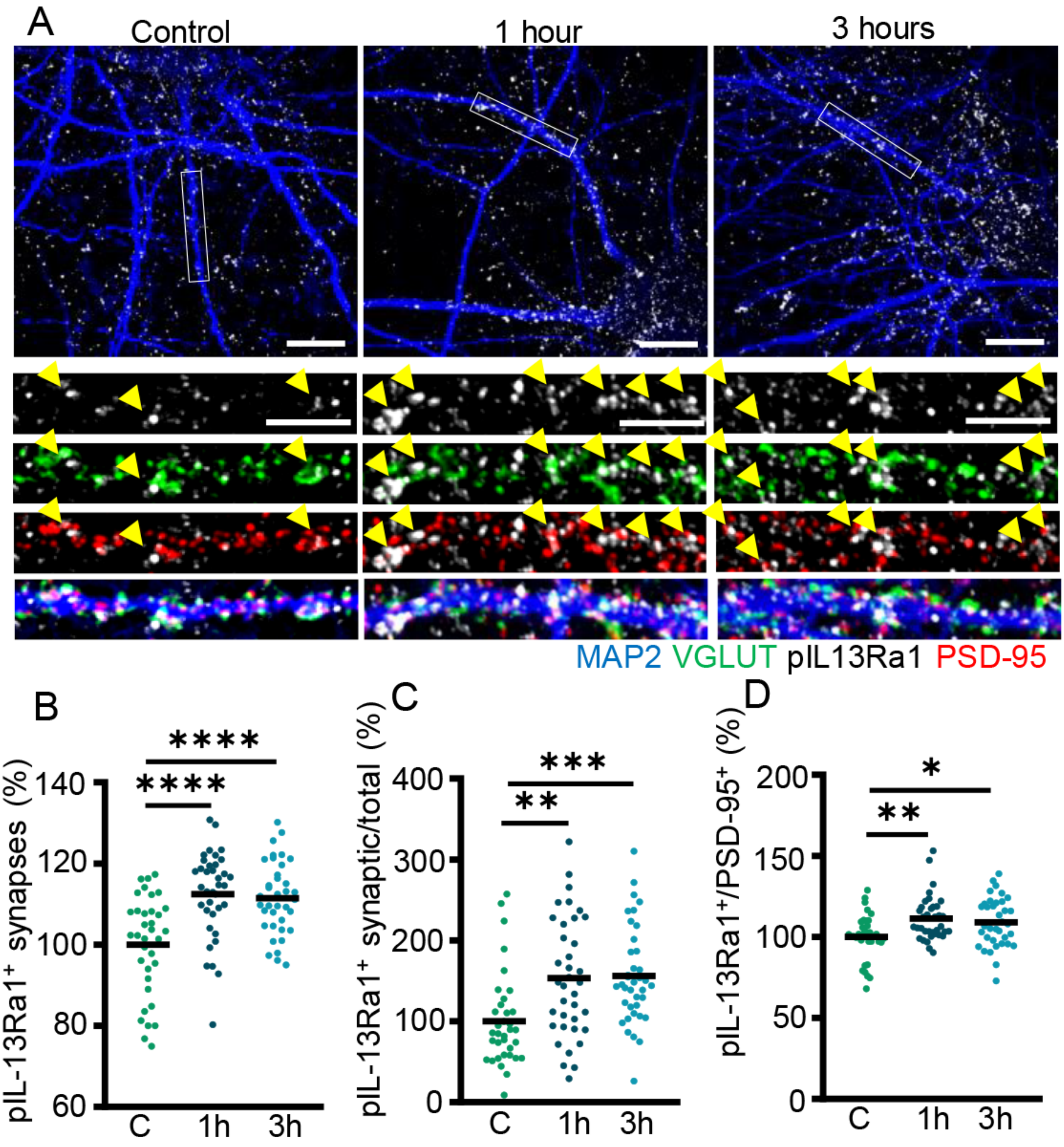

### supplementary figure 5

Supplementary_Figure_5.pdf
1610 x 1314

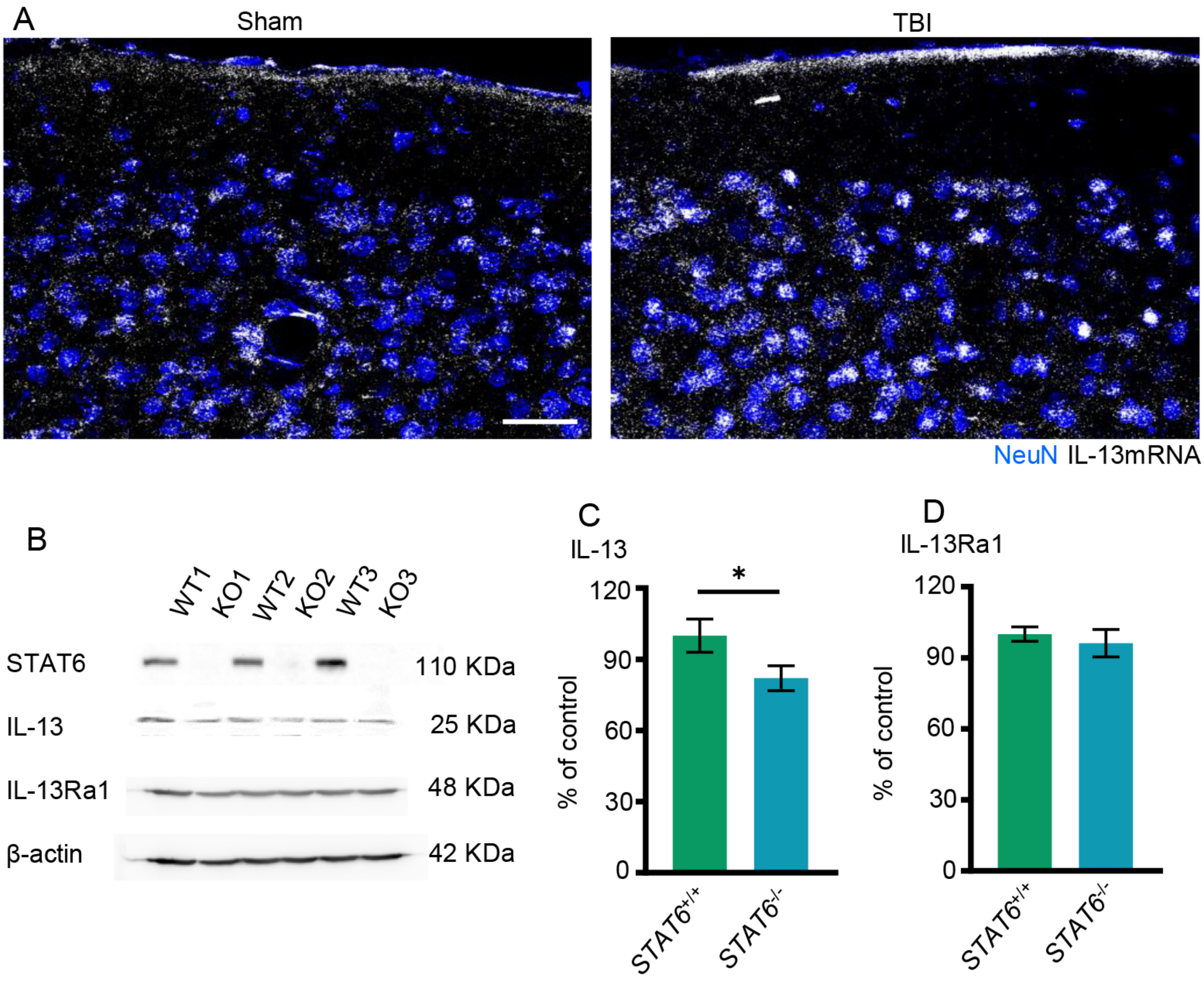

### supplementary figure 6

Supplementary_Figure_6.pdf

1266 x 1458

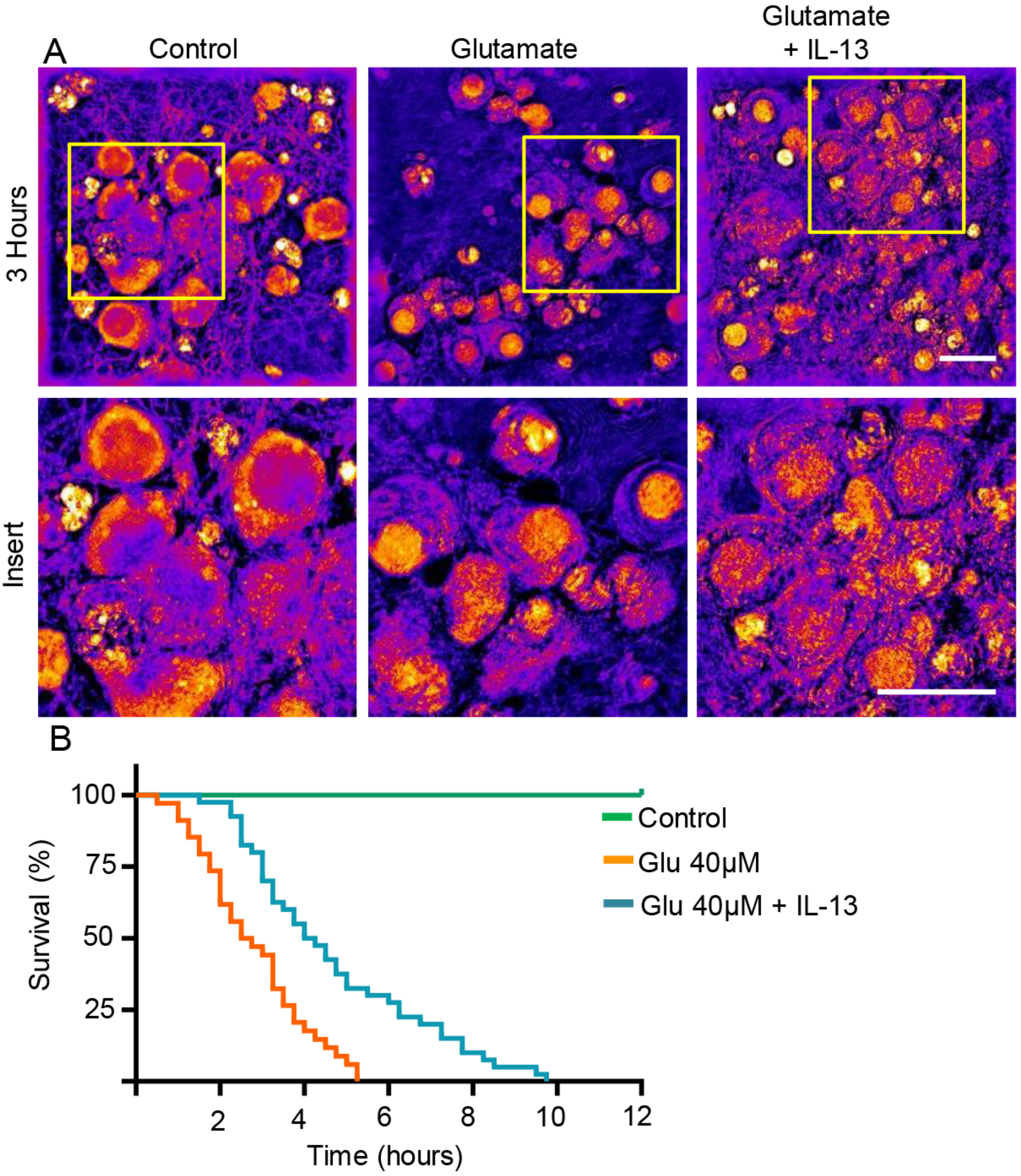
