## supplementary table 1 for "Interleukin-13 and its receptor are synaptic proteins involved in plasticity and neuroprotection"

| Supplementary table 1: List of antibodies used for immunostaining (IF), immunohistochemistry (IHC), Fluorescent in Situ Hybridisation (RNAscope) or for western blot (WB). | | | | |
| --- | --- | --- | --- | --- |
| **Primary antibodies** | **Host** | **Company** | **Catalog nr.** | **Dilution (application** |
| ATF-3 | Rabbit | Sigma | HPA001562 | 1:1000 (IF) |
| β-actin | Mouse | Proteintech | 69009-I-Ig | 1:10000 (WB) |
| Bassoon | Mouse | Enzo | SAP7F407 | 1:400 (IF) |
| c-fos | Mouse | Abcam | ab208942 | 1:500 (IF) |
| CREB(phospho) | Rabbit | CST | 9198S | 1:500 (IF) |
| ERK1/2(phospho) | Rabbit | CST | 9101 | 1:200 (IF) |
| ERK1/2(phospho) | Rabbit | CST | 4370 | 1:200 (IF) |
| GFP | Chicken | Abcam | ab13970 | 1:500 (RNAscope) |
| GFAP | Mouse | Sigma | G3893 | 1:500 (IF) |
| Homer-1 b/c | Rabbit | SySy | 160022 | 1:500 (IF) |
| Iba1 | Rabbit | SySy | 234003 | 1:1000 (IF) |
| IL-13 | Mouse | Santa | SC-393365 | 1:50 (IF), 1:100 (WB), 1:200 (IHC) |
| IL-13 | Rabbit | Bioss | bs-0560R | 1:100 (IF) |
| IL-13Rα1 | Rabbit | Invitrogen | PA5-50989 | 1:500 (IF, WB), 1:200 (IHC) |
| IL-13Rα1 (phospho) | Rabbit | Invitrogen | PA5-38607 | 1:250 (IF) |
| KCHiP3 (Dream) | Rabbit | Thermo | NPA5-11665 | 1:25 (IF) |
| Map2 | Chicken | Encor | CPCA-MAP2 | 1:1000 (IF) |
| Neu-N | Guinea pig | SySy | 266004 | 1:500 (IF), 1:200 (RNAscope) |
| PSD-95 | Mouse | Abcam | ab2723 | 1:500 (IF), 1:4000 (WB) |
| RFP (conjugated) | Camel | Nanotech | N0404-AT565-L | 1:500 (IF) |
| Shank2 | Rabbit | Homemade | SA5192 | 1:500 (IF) |
| STAT3(phospho) | Mouse | CST | 9138S | 1:200 (IF) |
| STAT6(phospho) | Rabbit | Invitrogen | PA5-104692 | 1:250 (IF) |
| STAT6 | Rabbit | CST | 9362 | 1:1000 (WB) |
| Synaptotagmin | Rabbit | SySy | 105 103C3 | 1:500 (IF) |
| Synaptophysin | Guinea pig | SySy | 101004 | 1:500 (IF) |
| Synaptophysin | Rabbit | Abcam | ab14692 | 1:100 (IF), 1:20000 (WB) |
| VGAT | Guinea pig | SySy | 131004 | 1:500 (IF) |
| VGLUT1 | Guinea pig | SySy | 135304 | 1:500 (IF) |
| VGLUT2 | Guinea pig | SySy | 135404 | 1:500 (IF) |

| **secondary antibodies** | **Host** | **Company** | **Catalog nr.** | **Dilution (application)** |
| --- | --- | --- | --- | --- |
| Anti-chicken AF 405 | Goat | Abcam | ab175674 | 1:500 (IF) |
| Anti-chicken AF 488 | Donkey | Biotium | 20166 | 1:500 (IF, RNAscope) |
| Anti-guinea pig AF 488 | Goat | Invitrogen | A11073 | 1:500 (IF) |
| Anti-guinea pig CF 568 | Donkey | Biotium | 20377 | 1:500 (IF) |
| Anti-guinea pig AF 633 | Donkey | Invitrogen | A21105 | 1:500 (RNAscope) |
| Anti-rabbit AF 488 | Donkey | Invitrogen | A21206 | 1:500 (IF) |
| Anti-rabbit AF 568 | Donkey | Invitrogen | A10042 | 1:500 (IF) |
| Anti-rabbit AF 647 | Donkey | Invitrogen | A31573 | 1:500 (IF) |
| Anti-mouse AF 488 | Donkey | Invitrogen | A21202 | 1:500 (IF) |
| Anti-mouse AF 647 | Donkey | Invitrogen | A31571 | 1:500 (IF) |
| DAPI 405 | - | Thermo | 62247 | 1:1000 (IF) |
| FluoTag-X4 anti-Rabbit AberriorStar 580 | - | Nanotag | N2404-Ab580-S | 1:500 (IF, STED) |
| FluoTag-X2 anti-Mouse AberriorStar635 | - | Nanotag | N1202-Ab635P-S | 1:500 (IF; STED) |
| Anti-mouse | Horse | Vector Lab | PI-2000 | 1:3000 (WB), 1:200 (IHC) |
| Anti-rabbit | Goat | Vector Lab | PI-1000 | 1:5000 (WB), 1:200 (IHC) |
