## supplementary table 2 for "Interleukin-13 and its receptor are synaptic proteins involved in plasticity and neuroprotection"

| **Gene (target)** | **Sequence** |
| --- | --- |
| *c-fos (rat)* | forward: 5’ -gagccgcgaacgagcagtga- 3’  Reverse: 5’ -ggcgaggggtccaggggtag- 3’ |
| *egr-1 (rat)* | forward: 5’ -aacaaccctacgagcacctg- 3’  Reverse: 5’ -accagcgccttctcgttatt- 3’ |
| *egr-2 (rat)* | forward: 5’ -aggccgtagacaaaatcccag- 3’  Reverse: 5’ -cctcccagttcaccattggg- 3’ |
| *fos-B (rat)* | forward: 5’ -gtgagagatttgccagggtc- 3’  Reverse: 5’ -agagagaagccgtcaggttg- 3’ |
| *gadd45a (rat)* | forward: 5’ -ggagtcagcgcaccataact- 3’  Reverse: 5’ -ggtcgtcatcttcatccgca- 3’ |
| *gadd45b (rat)* | forward: 5’ -ctcctggtcacgaactgtcat- 3’  Reverse: 5’ -ggacccactggttattgcct- 3’ |
| *GAPDH (rat)* | forward: 5’ -gacatgccgcctggagaaac- 3’  Reverse: 5’ -agcccaggatgccctttagt- 3’ |
| *IL-13 (rat)* | forward: 5’ -atggtatggagcgtggacct- 3’  Reverse: 5’ -actggagatgttggtcaggg- 3’ |
| *IL-13Ra1 (rat)* | forward: 5’ -tgagtctgctgtgaccgaac- 3’  Reverse: 5’ -atagttggtgtccgggcttg- 3’ |
| *IL-13Ra2 (rat)* | forward: 5’ -cacagggccagactcaaagat- 3’  Reverse: 5’ -gtgggttcagggtcttccttt- 3’ |
| *BDNF (human)* | forward: 5’ -catccgaggacaaggtggcttg- 3’  Reverse: 5’ -gccgaactttctggtcctcatc- 3’ |
| *GAPDH (human)* | forward: 5’ -gtctcctctgacttcaacagcg- 3’  Reverse: 5’ -accaccctgttgctgtagccaa- 3’ |
| *IL-13 (human)* | forward: 5’ -atgcatccgctcctcaatcc- 3’  Reverse: 5’ -agtgagagcaatgaccgtgg- 3’ |
| *IL-13Ra1 (human)* | forward: 5’ -cctgaatgagaggatttgtctgc- 3’  Reverse: 5’ -cagtcacagcagactcaggatc- 3’ |
| *IL-13Ra2 (human)* | forward: 5’ -gtggagtgataaacaatgctggg- 3’  Reverse: 5’ -tgggtaggtgtttggcttacgc- 3’ |
| *TNF-α (human)* | forward: 5’ -ctcttctgcctgctgcactttg- 3’  Reverse: 5’ -atgggctacaggcttgtcactc- 3’ |

Supplementary Table 2: List of genes and sequences used for RT-qPCR.
